# Identification of a series of pyrrolo-pyrimidine based SARS-CoV-2 Mac1 inhibitors that repress coronavirus replication

**DOI:** 10.1101/2024.10.28.620664

**Authors:** Jessica J. Pfannenstiel, Men Thi Hoai Duong, Daniel Cluff, Lavania M. Sherrill, Iain Colquhoun, Gabrielle Cadoux, Devyn Thorne, Johan Pääkkönen, Nathaniel F. Schemmel, Joseph O’Connor, Pradtahna Saenjamsai, Mei Feng, Michael J. Hageman, David K. Johnson, Anuradha Roy, Lari Lehtiö, Dana V. Ferraris, Anthony R. Fehr

**Author notes:** Corresponding authors: *Email addresses:* (L. Lehtiö), (D. Ferraris), (A.R. Fehr).

## Abstract

Coronaviruses (CoVs) can emerge from zoonotic sources and cause severe diseases in humans and animals. All CoVs encode for a macrodomain (Mac1) that binds to and removes ADP-ribose from target proteins. SARS-CoV-2 Mac1 promotes virus replication in the presence of interferon (IFN) and blocks the production of IFN, though the mechanisms by which it mediates these functions remain unknown. Mac1 inhibitors could help elucidate these mechanisms and serve as therapeutic agents against CoV-induced diseases. We previously identified compound **4a** (a.k.a. MCD-628), a pyrrolo-pyrimidine that inhibited Mac1 activity *in vitro* at low micromolar levels. Here, we determined the binding mode of **4a** by crystallography, further defining its interaction with Mac1. However, **4a** did not reduce CoV replication, which we hypothesized was due to its acidic side chain limiting permeability. To test this hypothesis, we developed several hydrophobic derivatives of **4a**. We identified four compounds that both inhibited Mac1 *in vitro* and inhibited murine hepatitis virus (MHV) replication: **5a**, **5c**, **6d**, and **6e**. Furthermore, **5c** and **6e** inhibited SARS-CoV-2 replication only in the presence of IFN*γ*, similar to a Mac1 deletion virus. To confirm their specificity, we passaged MHV in the presence of **5a** to identify drug-resistant mutations and identified an alanine-to-threonine and glycine-to-valine double mutation in Mac1. Recombinant virus with these mutations had enhanced replication compared to WT virus when treated with **5a**, demonstrating the specificity of these compounds during infection. However, this virus is highly attenuated *in vivo*, indicating that drug-resistance emerged at the expense of viral fitness.

**IMPORTANCE:** Coronaviruses (CoVs) present significant threats to human and animal health, as evidenced by recent outbreaks of MERS-CoV and SARS-CoV-2. All CoVs encode for a highly conserved macrodomain protein (Mac1) that binds to and removes ADP-ribose from proteins, which promotes virus replication and blocks IFN production, though the exact mechanisms remain unclear. Inhibiting Mac1 could provide valuable insights into these mechanisms and offer new therapeutic avenues for CoV-induced diseases. We have identified several unique pyrrolo-pyrimidine-based compounds as Mac1 inhibitors. Notably, at least two of these compounds inhibited both murine hepatitis virus (MHV) and SARS-CoV-2 replication. Furthermore, we identified a drug-resistant mutation in Mac1, confirming target specificity during infection. However, this mutant is highly attenuated in mice, indicating that drug-resistance appears to come at a fitness cost. These results emphasize the potential of Mac1 as a drug target and the promise of structure-based inhibitor design in combating coronavirus infections.

## INTRODUCTION

Coronaviruses (CoVs) are large, positive-sense RNA viruses that infect a wide variety of mammalian species, including humans. Some human CoVs (HCoVs), such as HCoV-OC43, HKU1, NL63, and HCoV-229E are endemic and contribute to the common cold, while others, severe acute respiratory syndrome (SARS)-CoV, Middle East respiratory syndrome (MERS)-CoV, and SARS-CoV-2 have caused epidemic outbreaks of severe disease and human fatalities. The recent COVID-19 pandemic caused by SARS-CoV-2 resulted in the deaths of over seven million people worldwide and had profound social and economic consequences. Beyond the devastating human toll, the pandemic disrupted healthcare systems, led to widespread economic downturns, and prompted significant changes in global public health infrastructure and policy (1). SARS-CoV-2 is now endemic in the human population and continues to cause severe disease in humans. Furthermore, many other CoVs have been identified in wildlife, posing a continuous threat of zoonotic transmission that could lead to additional epidemics (2). Thus, there is an urgent need for novel therapeutic interventions and further vaccine development.

Coronaviruses (CoVs) evade the host’s innate immune response by encoding for multiple proteins that either repress the production of interferon (IFN) or directly inhibit IFN-stimulated genes (ISGs) (3, 4). Several PARP proteins are highly induced by IFN and are part of the antiviral response (5, 6). Most PARPs act as ADP-ribosyltransferases (ARTs) that add single (mono) or multiple (poly) units of ADP-ribose onto proteins (7). ADP-ribosylation can be reversed by several different classes of enzymes, including macrodomains (8). All CoVs, alphaviruses, Hepatitis E virus, and Rubella virus encode a macrodomain in their genome, indicating that a broad spectrum of positive-sense RNA viruses utilize macrodomains to reverse ADP-ribosylation during infection (9, 10). For CoVs, the conserved macrodomain is encoded within non-structural protein 3 (nsp3), and is called Mac1. Mac1 binds to and removes ADP-ribose from protein, countering host PARPs (6, 11). Prior research has shown that murine hepatitis virus (MHV), SARS-CoV, MERS-CoV, and SARS-CoV-2 viruses engineered with point mutations that reduce Mac1 ADP-ribose binding or hydrolysis activity replicate poorly, lead to enhanced IFN and pro-inflammatory cytokine responses and cause minimal disease in animal models of infection (12-20). Furthermore, recombinant alphaviruses with macrodomain mutations are also highly attenuated in cell culture and in mice (21-23). Understanding the role of viral macrodomains in immune evasion and viral replication is crucial for developing effective therapeutic interventions and vaccines.

Interestingly, the complete deletion of SARS-CoV-2 Mac1 does not substantially impair viral replication in cell culture, which contrasts with other CoVs, such as MHV and MERS-CoV, where Mac1 deletion leads to unrecoverable viruses (18). However, SARS-CoV-2 Mac1-deletion and point mutant viruses exhibited increased sensitivity to IFN-*γ*, led to increased production of IFN and ISGs, and did not cause severe disease in mice (18-20). This indicates that SARS-CoV-2 Mac1 plays a critical role during infection, though its specific targets during infection and downstream consequences, such as its effect on the viral lifecycle, remain largely unknown. Mac1 inhibitors could thus be useful tools to help identify these targets and better understand how Mac1 directly promotes virus replication and pathogenesis. Furthermore, as Mac1 is completely conserved across all CoVs and is vital for viral pathogenesis, it could be a unique therapeutic target for SARS-CoV-2 and other potential pandemic CoVs (2).

Since the outbreak of SARS-CoV in 2003 and up to the start of the COVID-19 pandemic, several studies have determined the structure of Mac1 from multiple CoVs and alphaviruses, including SARS-CoV, 229E, Infectious Bronchitis Virus (IBV), MERS-CoV, HKU4, Chikungunya virus (CHIKV) and Venezuelan Equine Encephalitis virus (VEEV) (24-29). Much like macrodomains that had been discovered from species such as archaea (30), the viral macrodomains form an *αβα* sandwich like structure with several *β*-sheets surrounded by *α*-helices on both sides, with a highly defined ADP-ribose binding pocket. Shortly after the pandemic began, several SARS-CoV-2 Mac1 structures were determined, which provided detailed atomic-level resolution of the SARS-CoV-2 Mac1 protein facilitating drug-discovery efforts (11, 31-34).

Multiple groups have now identified Mac1 inhibitors through high-throughput screening and targeted drug development using these crystal structures (35-44). Through these efforts, several compounds with IC_50_ values between 0.4 and 10 *μ*M have been discovered with high specificity *in vitro* (45). One of these studies utilized a unique crystallography-based fragment screen that identified several small molecules that bound to the ADP-ribose binding pocket of Mac1 (35). These fragments served as promising starting points for further inhibitor development for multiple groups (36, 37, 39). Starting with a small pyrrolo-pyrimidine fragment with weak *in vitro* potency (IC_50_ of 180 *μ*M), we synthesized a series of primary and secondary amino acid-based pyrrolo-pyrimidines to determine whether more potent Mac1 inhibitors could be developed. The previously described luminescent-based AlphaScreen™ (AS) assay was utilized to screen ∼60 pyrrolo-pyrimidines for their ability to inhibit Mac1-ADP-ribose binding. Of these pyrrolo pyrimidines, we identified a tryptophanate (MCD-628) that inhibited SARS-CoV-2 Mac1-ADP-ribose binding with an IC_50_ of 6.1 *μ*M. MCD-628 incubation with Mac1 also increased its thermal stability to nearly the same degree as ADP-ribose, indicating that it directly binds to Mac1. However, this compound contains a carboxylic acid group that suggests it would have poor permeability and be unlikely to inhibit virus replication. Many potent Mac1 inhibitors have acidic or highly polar moieties, likely limiting their ability to inhibit virus replication or pathogenesis (45). To date, only one Mac1 inhibitor has been published that represses CoV replication in cell culture (40). Thus, future Mac1 inhibitors must be designed to increase their inhibition of Mac1 biochemical functions *in vitro* and repress CoV replication in cell culture or animal model infections.

In this study, we solved a co-crystal structure of Mac1 with MCD-628 (termed **4a** herein) and designed a novel series of pyrrolo-pyrimidine-based Mac1 inhibitors with increased lipophilicity to identify compounds that could both inhibit Mac1 *in vitro* and repress CoV replication in cell culture. We replaced the acidic moiety of our lead compound, MCD-628 (termed **4a** herein), with several esters and amide couplings with hydrophobic pyridines. These modifications largely improved cell permeability while maintaining inhibitory activity *in vitro*. We identified four compounds that substantially repressed MHV replication in cell culture, including two that inhibited MHV and SARS-CoV-2. Importantly, these compounds only inhibited SARS-CoV-2 in the presence of IFN-*γ*, in-line with results demonstrating that Mac1-deleted or mutated SARS-CoV-2 viruses are highly sensitive to IFN-*γ* (18-20). Additionally, mutations conferring resistance to one of these inhibitors were identified, further confirming their target specificity. These findings demonstrate that Mac1 inhibitors can repress virus replication and offer a promising platform for developing Mac1 chemical probes and CoV antivirals.

## RESULTS

### Structural and biochemical analysis of pyrrolo-pyrimidine based SARS-CoV-2 Mac1 inhibitor

Using a series of amino acid based 7H-pyrrolo[2,3-d] pyrimidines, we previously created compound **4a** (*S*), derived from tryptophan, that inhibited SARS-CoV-2 Mac1 binding to ADP-ribose with a 6.1 *μ*M IC_50_ value (Fig. 1A-B) (39). **4a** also inhibited Mac1 enzyme activity and bound to Mac1 as demonstrated by a thermal shift profile similar to Mac1’s natural ligand, ADP-ribose (39). Having established that **4a** binds and inhibits Mac1 *in vitro*, we sought to get more insight into the mechanism by which **4a** binds to Mac1 by determining the structure of **4a** with Mac1. We solved the structure of **4a** with Mac1 and refined it to 1.1 Å resolution (Fig. 1C-D, Table S1). In this structure, some of the key features are a hydrogen bond with D22 and the backbone of I23, multiple hydrogen bonds between the carboxylate with neighboring water molecules, and finally a hydrogen bond between the indole NH and L126 (Fig. 1C-D). F156, previously seen to form pi stacking interactions with ADP-ribose and inhibitors, has considerable flexibility, and while it is next to the pyrrolo-pyrimidine and contributes to hydrophobic interactions, the geometry does not allow pi-stacking interactions. This co-crystal structure provides a strong starting point for the synthesis of additional Mac1 inhibitors.

**Fig 1.**
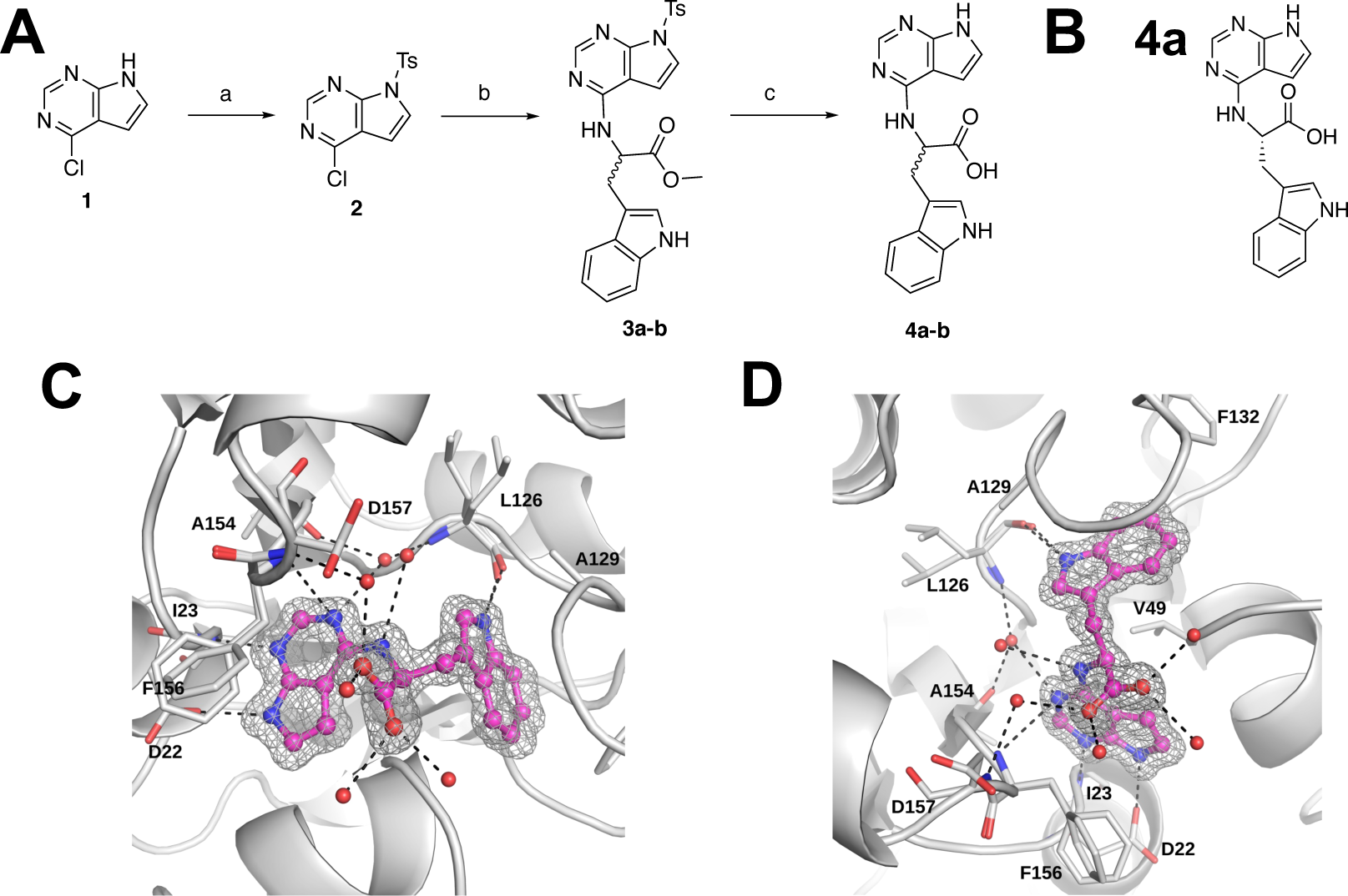
Crystal structure of 4a provides new insight into its interaction with Mac1. (A-B) Chemical synthesis plan to produce **4a-b** (A), and the chemical structure of **4a** (B). (C-D) Crystal structure of **4a** in two different poses (PDB id. 9GUB). These poses include images where the pyrrolo-pyrimidine is oriented in the front left (C) or in the lower middle (D). Note that the carboxylate makes hydrogen bonds with 3 different water molecules and the tryptophanate makes a hydrogen bond with the backbone of L126. The sigma-A weighted 2Fo-Fc electron density map is contoured at 1.0 σ. Waters are shown as red spheres, and hydrogen bonds are illustrated as black dashed lines.

Next, we tested whether the enantiomer of **4a**, **4b** (*R*) (Fig. 2A), would also bind and inhibit SARS-CoV-2 Mac1. Indeed, **4b** interacted with Mac1 as it had a similar thermal shift profile to that of **4a** (Fig. 2B). Furthermore, **4b** inhibited Mac1 binding to an ADP-ribosylated peptide in an AlphaScreen assay with an IC_50_ of 1.66 *μ*M with almost no inhibition of the Bn-His peptide control (Fig. 2C). We were able to reproduce the experimental binding mode of 4a by molecular modeling and subsequently demonstrated that **4b** would interact with Mac1 in a similar manner (Fig. 2D).

**Fig 2.**
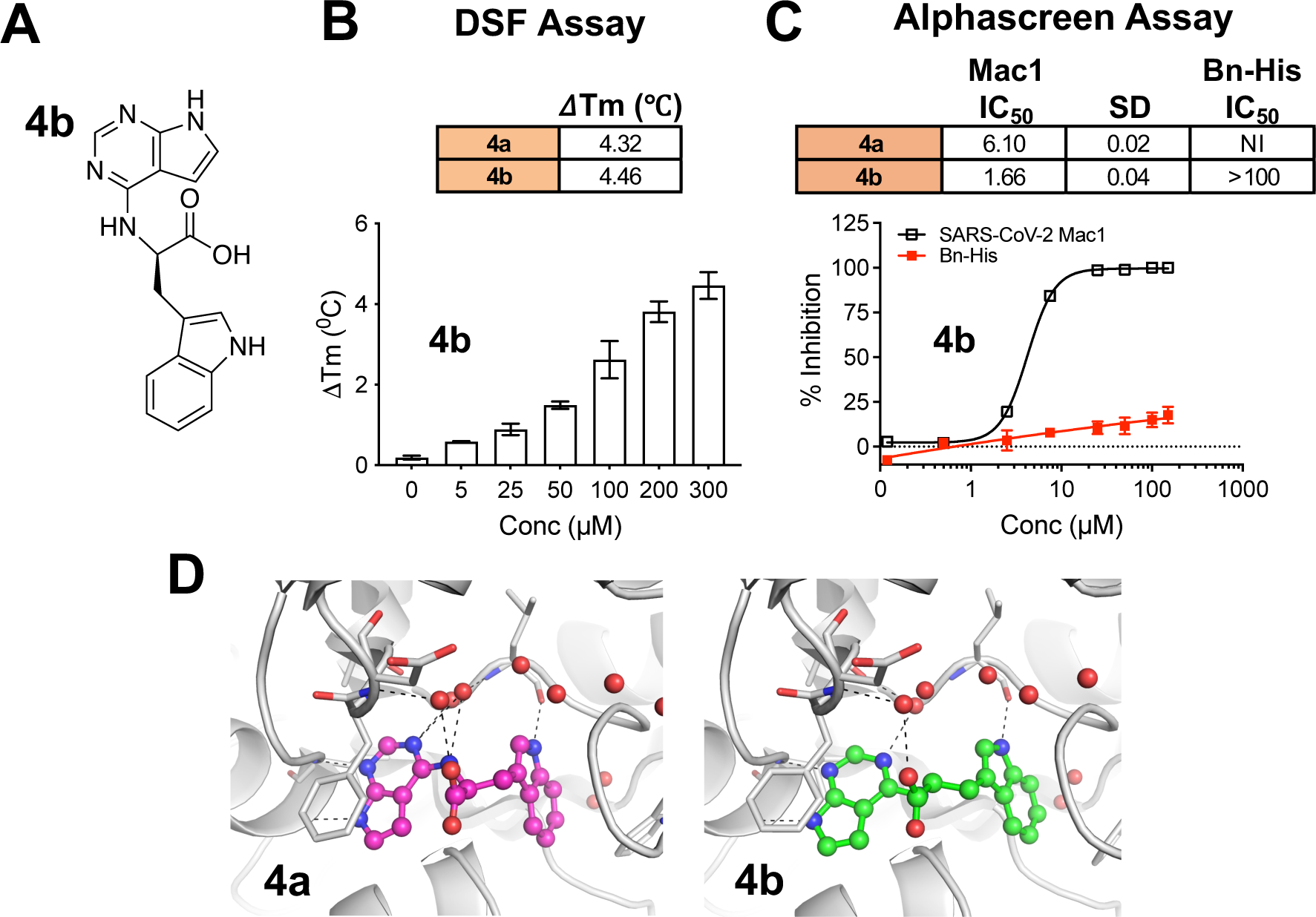
Compound 4b interacts with Mac1 and inhibits Mac1-ADP-ribose binding. A) Chemical structure of compound **4b**. B) Compound **4b** was incubated with SARSCoV-2 Mac1 at increasing concentrations and the thermal stability of SARS-CoV-2 Mac1 was determined by a DSF assay. The *Δ*Tm is the average 5 experimental replicates. n=5. C) Competition assays were used to demonstrate that **4a** and **4b** block the interaction between Mac1 and ADP-ribosylated peptides in the AS assay. The IC50 represents the average value of 2 independent experiments, each done with 3 experimental replicates. The graphs are from one experiment representative of 2 independent experiments. D) Compounds **4a** and **4b** were docked into Mac1 using PDB: 9GUB. Hydrogen bonds are illustrated as dashed lines.

Compounds **4a**/**4b** have a negatively charged carboxylic acid moiety at physiological pH, which we hypothesized might preclude its ability to cross cellular membranes. To address this potential problem, we first replaced the carboxylic acid with methyl (**5a**/**5b**) and isopropyl (**5c**/**5d**) esters (Fig 3A). Unexpectantly, the esters derived from **4b**, **5b** and **5d**, did not demonstrate any significant inhibition of Mac1 in the AlphaScreen assay (data not shown), while the **4a** derivatives, **5a** and **5c**, interacted with Mac1 by the thermal shift assay (Fig. 3B) and inhibited Mac1 in the AlphaScreen assay with IC_50_ values of 14.14 *μ*M and 3.66 *μ*M, respectively (Fig. 3C). Based on modeling, the additional carbon atoms on these molecules appear to protrude out of the binding pocket and have only minimal impact on the overall interaction of these compounds with Mac1 (Fig. 3D). Importantly, the addition of the esters dramatically increased the lipophilicity of these compounds, as the logD at pH 7.4 of these compounds went from -0.61 (**4a**) to 1.5 (**5a**) and 3.33 (**5c**), indicating that the ester-modified compounds are much more likely to cross cellular membranes and target Mac1 during infection.

**Fig 3.**
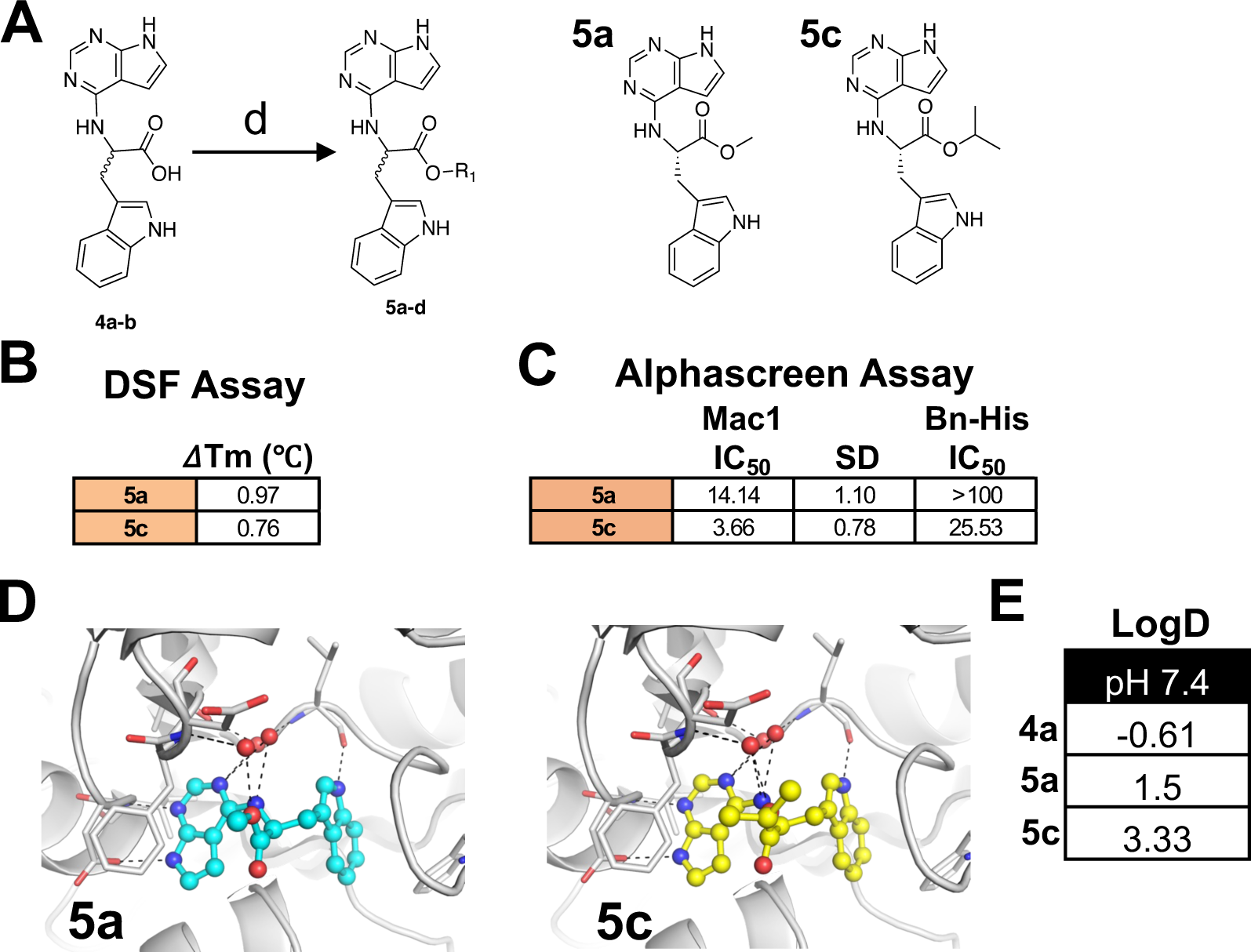
Compounds 5a and 5c interact with Mac1 and inhibit Mac1-ADP-ribose binding. A) Modification of **4a-b** to ester derivatives **5a** and **5c**. B) Compounds **5a** and **5c** were incubated with SARS-CoV-2 Mac1 at increasing concentrations and the thermal stability of SARS-CoV-2 Mac1 was determined by a DSF assay. The *Δ*Tm is the average 5 experimental replicates. n=5. C) Competition assays were used to demonstrate that **5a** and **5c** block the interaction between Mac1 and ADP-ribosylated peptides in the AS assay. The IC_50_ represents the average value of 2 independent experiments. n=3 experimental replicates. D) Compounds **5a** and **5c** were docked into Mac1 using PDB: 9GUB. Hydrogen bonds are illustrated as dashed lines. E) The LogD values were experimentally determined using the shake-flask method for **4a**, **5a**, and **5c**.

### Pyrrolo-pyrimidine based esters inhibits MHV-JHM replication

We next aimed to determine if these compounds could inhibit CoV replication. A SARS-CoV-2 Mac1-deletion virus had only a modest growth defect in Calu-3 cells of 2-3-fold, which indicates that Mac1 inhibitors may not impact SARS-CoV-2 replication in cell culture. In contrast, Mac1 is critical, if not essential, for the replication of murine hepatitis virus (MHV) strain JHM (JHMV) (17). Thus, we hypothesized that this virus may be better suited for testing Mac1 inhibitors for their ability to inhibit virus replication. To enable more efficient screening of compounds for impacts on virus replication, we replaced ORF4 of JHMV with nanoluciferase (JHMV-nluc) (Fig. S1), as the deletion of ORF4 does not affect JHMV replication or pathogenesis (46). JHMV-nluc replicated like WT virus (Fig. 4A) and importantly, produced over 10^6^ light units at peak replication (Fig. 4B). Next, we tested the ability of **4a**, **4b**, **5a**, and **5c** to inhibit JHMV-nluc replication in DBT cells at concentrations ranging from 25 to 200 *μ*M, using **GS441524** (active metabolite of remdesivir) as a control. DBT cells are an astrocytoma cell line that are susceptible to MHV. JHMV replication in DBT cells is highly dependent on Mac1 activity, as a D1329A mutant virus replicates very poorly in these cells (17). As expected, **4a** and **4b** did not affect JHMV replication, as opposed to **GS441524**, which inhibited virus replication at all concentrations. In contrast, both **5a** and **5c** inhibited JHMV replication, with **5c** being significantly more potent, having inhibited JHMV to nearly the same level as **GS441524** at 25 *μ*M (Fig. 4C). Importantly, none of these molecules showed substantial cytotoxicity at the concentrations tested (Fig. S2A-B).

**Fig 4.**
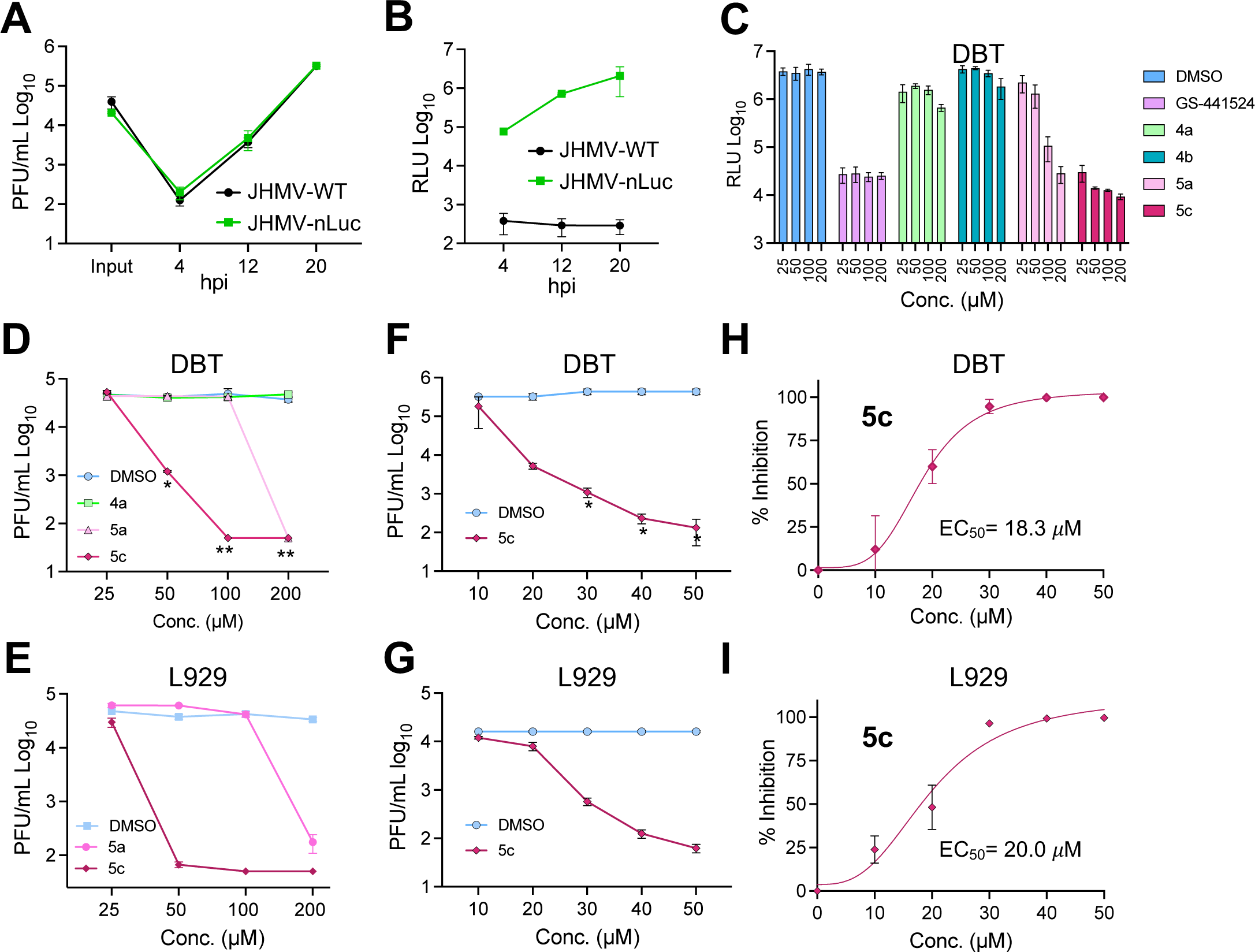
Compounds 5a and 5c, but not 4a, inhibit MHV replication. A) 17Cl-1 cells were infected with JHMV-WT and JHMV-nLuc viruses at an MOI = 0.1. Cells and supernatants were collected at indicated timepoints and progeny virus was determined by plaque assay. B) 17Cl-1 cells were infected as described in A. Lysates were collected at indicated times and luciferase activity was determined using a nano-Glo luciferase assay kit measured as per manufacturer’s instructions. The results in A and B are from 1 experiment representative of 2 independent experiments. N=3 biological replicates. C) DBT cells were infected with JHMV-nLuc at an MOI = 0.1 and at 1 hpi, the indicated concentration of each compound was added to the media. Lysates were collected at 20 hpi and luciferase activity was measured as described in B. D-E) DBT cells were infected with JHMV-WT and at 1 hpi the indicated concentration of each compound was added to the media. Cells and supernatants were collected at 20 hpi and progeny virus was measured by plaque assay. The results in C-E are from 1 experiment representative of 2 independent experiments. n=3 biological replicates. F) The combined average % JHMV-WT inhibition of 10-50 *μ*M **5c** on DBT cells over 2 independent experiments. G-H) L929 cells were infected with JHMV-WT and at 1 hpi the indicated concentration of each compound was added to the media. Cells and supernatants were collected at 20 hpi and progeny virus was measured by plaque assay. The results in G-H are from 1 experiment representative of 2 independent experiments. n=3. I) The combined average % JHMV-WT inhibition of 10-50 *μ*M **5c** on DBT cells over 3 independent experiments. The results in C, D, E, G, and H are from 1 experiment representative of 3 independent experiments. N=3 biological replicates.

Next, we tested whether **5a** or **5c** would impact the production of infectious virus. Indeed, we found that both **5a** and **5c** inhibited the production of infectious virus following infection of both DBT and L929 cells with JHMV (Fig. 4D-E). **5a** only inhibited virus production at 200 *μ*M, while **5c** inhibited virus replication with as little as 25 *μ*M and decreased replication by ∼1.5 and 3 logs at 50 *μ*M on DBT and L929 cells, respectively. To better determine the EC_50_ for **5c**, we tested its activity at concentrations from 0-50 *μ*M on DBT, L292, and 17Cl-1 cells (Fig. 4F-G, S3A). The inhibition of virus production at these concentrations was dose-dependent and using this data we determined that the EC_50_ for **5c** on was ∼10-20 *μ*M, not substantially different from its IC_50_ of 3.66 *μ*M (Fig. 4H-I, S3B).

Next, amide couplings were conducted with carboxylates **4a** and **4b** to create 25 additional compounds, many of which included highly hydrophobic side chains to increase the lipophilicity. Of these, 5 compounds demonstrated IC_50_ values of less than 10 *μ*M in our initial screening and were named **6a**-**6e** (Fig. 5A and data not shown). Compounds **6a** and **6b** contain a pyridine group attached to the amide with the only difference being a chlorine atom on **6b**. **6d** *(S)* and **6e** *(R)* are enantiomers and only differ from **6a** in the position of the nitrogen on the pyridine. Finally, **6c** only has an amide group to replace the carboxylate. Only **6e** was derived from **4b**, while the rest were derived from **4a**. Following dose-response curves, we found that each of these 5 compounds had very similar IC_50_ values ranging from 4.0-8.4 *μ*M in the AlphaScreen assay (Fig. 5B). They also had thermal shifts of 1-3℃ in the DSF assay when incubated with the SARS-CoV-2 Mac1 protein, indicating a direct interaction with Mac1 (Fig. 5C). All of them had cLogD values between 1 and 3, indicating an increased lipophilicity compared to the parent compounds (**4a** LogD -0.61). Based on our modeling data, the position of each of these molecules in the binding pocket does not change significantly. The only major difference is the position of the pyridine for each molecule (Fig. 5D). For **6a**, **6b**, and **6d**, the pyridine protrudes out from the pocket and into slightly different poses for each one. In contrast, the pyridine of **6e** extends into the oxyanion subsite, which could explain its slightly greater inhibition of Mac1-ADP-ribose binding in the AlphaScreen assay.

**Fig 5.**
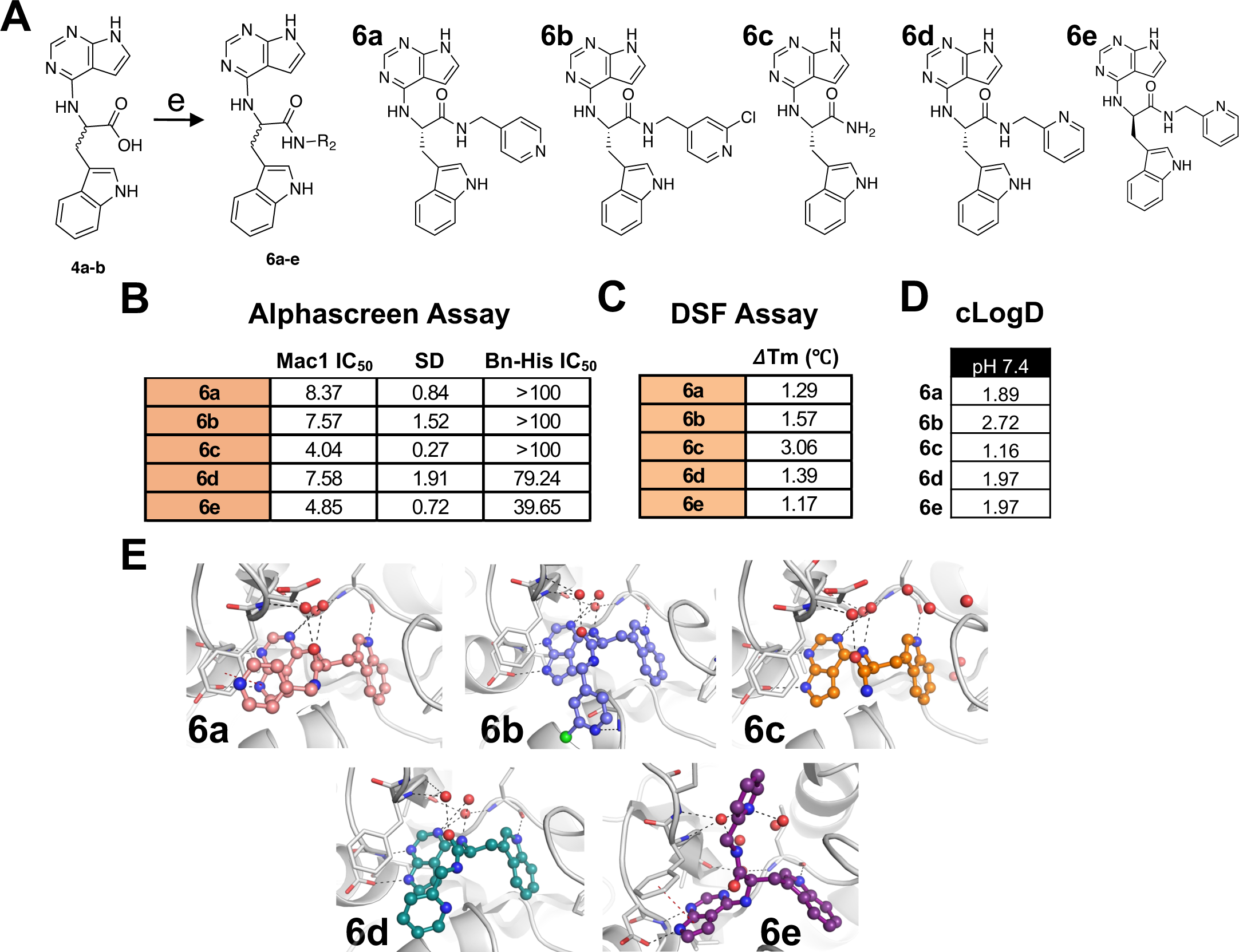
Group 6 compounds interact with Mac1 and inhibit Mac1-ADP-ribose binding. A) Modification of **4a-b** to several new derivatives, **6a-6e. 6a**-**6d** are derivatives of **4a** while **6e** is a derivates of **4b**. B) Competition assays were used to demonstrate that **6a-6e** block the interaction between Mac1 and ADP-ribosylated peptides in the AS assay. C) Compounds **6a-6e** were incubated with SARS-CoV-2 Mac1 at increasing concentrations and the thermal stability of SARS-CoV-2 Mac1 was determined by a DSF assay. Quantification data in B-C represents the average value of 2 independent experiments. D) Predicted cLogD values of **6a**-**6e**. E) Compounds **6a-6e** were docked into Mac1. Hydrogen bonds are illustrated as dashed lines.

### Inhibition of MHV-JHM replication by 6d and 6e

Similar to series **5**, we next tested if compounds in series **6** could inhibit virus replication. Using JHMV-nLuc, our initial screening found that only **6d** and **6e** inhibited light production from MHV replication in a dose-dependent manner (Fig. 6A). Furthermore, **6d** and **6e** demonstrated no substantial impact on cell viability (Fig. S4A-B). We next performed dose-response curves for **6d** and **6e** on both DBT and L929 cells at concentrations ranging from 50-200 *μ*M and found that the compounds inhibited MHV in a dose-dependent manner on both cells, with better activity on DBT cells (Fig. 6B-C). Using these results, we determined that the EC_50_ value for **6d** and **6e** on DBT cells was 77.1 and 53.5 *μ*M, respectively (Fig. 6D-E).

**Fig 6.**
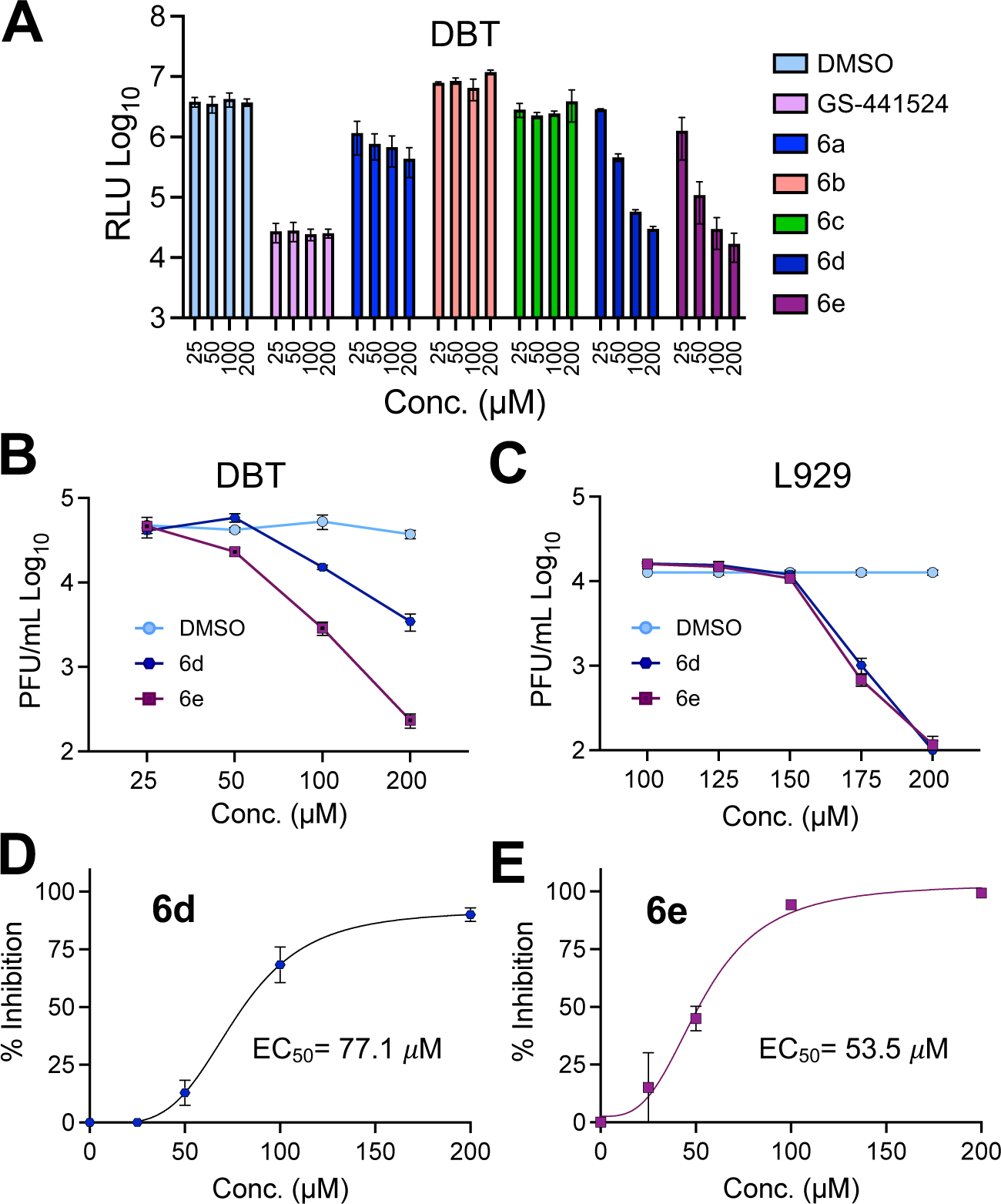
Group 6 compounds interact with Mac1 and inhibit Mac1-ADP-ribose binding. A) Modification of **4a-b** to several new derivatives, **6a-6e. 6a**-**6d** are derivatives of **4a** while **6e** is a derivates of **4b**. B) Competition assays were used to demonstrate that **6a-6e** block the interaction between Mac1 and ADP-ribosylated peptides in the AS assay. C) Compounds **6a-6e** were incubated with SARS-CoV-2 Mac1 at increasing concentrations and the thermal stability of SARS-CoV-2 Mac1 was determined by a DSF assay. The IC50 and *Δ*Tm values in B-C represent the average value of 2 independent experiments, each done with 3 experimental replicates. D) Predicted cLogD values of **6a**-**6e**. E) Compounds **6a-6e** were docked into Mac1 using PDB: 9GUB. Hydrogen bonds are illustrated as dashed lines.

### Compounds 5c and 6e inhibit SARS-CoV-2 replication

Having established antiviral activity against MHV, we next wanted to determine if our pyrrolo-pyrimidine based compounds could also inhibit SARS-CoV-2 replication. We hypothesized that our compounds would be more potent against SARS-CoV-2 as they were identified for their ability to inhibit the SARS-CoV-2 Mac1 protein, not the MHV Mac1 protein, *in vitro* (Figs. 3 & 5). Recently, we demonstrated that a full Mac1 deletion virus (SARS-CoV-2 *Δ*Mac1) replicates normally in cell culture compared to WT virus except when cells are pre-treated with IFN-*γ*. In the presence of 100 U of IFN-*γ Δ*Mac1 replicated ∼10-fold worse than WT virus in Calu-3 cells. Thus, we hypothesized that our compounds would only inhibit SARS-CoV-2 if cells are pre-treated with IFN-*γ*. So, we pretreated Calu-3 cells with IFN-*γ* and then infected cells in the presence or absence of **5c** or **6e** from 0 to 25 *μ*M (Fig. 7). First, we confirmed that neither **5c** nor **6e** affected the viability of Calu-3 cells (Fig. S5). In the absence of IFN-*γ*, **6e** did not reduce infectious virus production, and **5c** only reduced viral titers ∼2-fold at 25 *μ*M. In contrast, each compound reduced infectious virus production in the presence of IFN-*γ* at both 12.5 and 25 *μ*M (Fig. 7A-B). **5c** reduced viral titers by 3 and 7.5-fold, while **6e** reduced them by 2.7 and 3.8-fold at 12.5 and 25 *μ*M, respectively, indicating that the EC_50_ for each compound would be no greater than 12.5 *μ*M, again similar to their IC_50_ values. These results demonstrate that our pyrrolo-pyrimidine based Mac1 inhibitors are more potent against SARS-CoV-2 and the fact that they only inhibit virus production in the presence of IFN-*γ* strongly indicates that they specifically target Mac1 at the concentrations tested.

**Fig 7.**
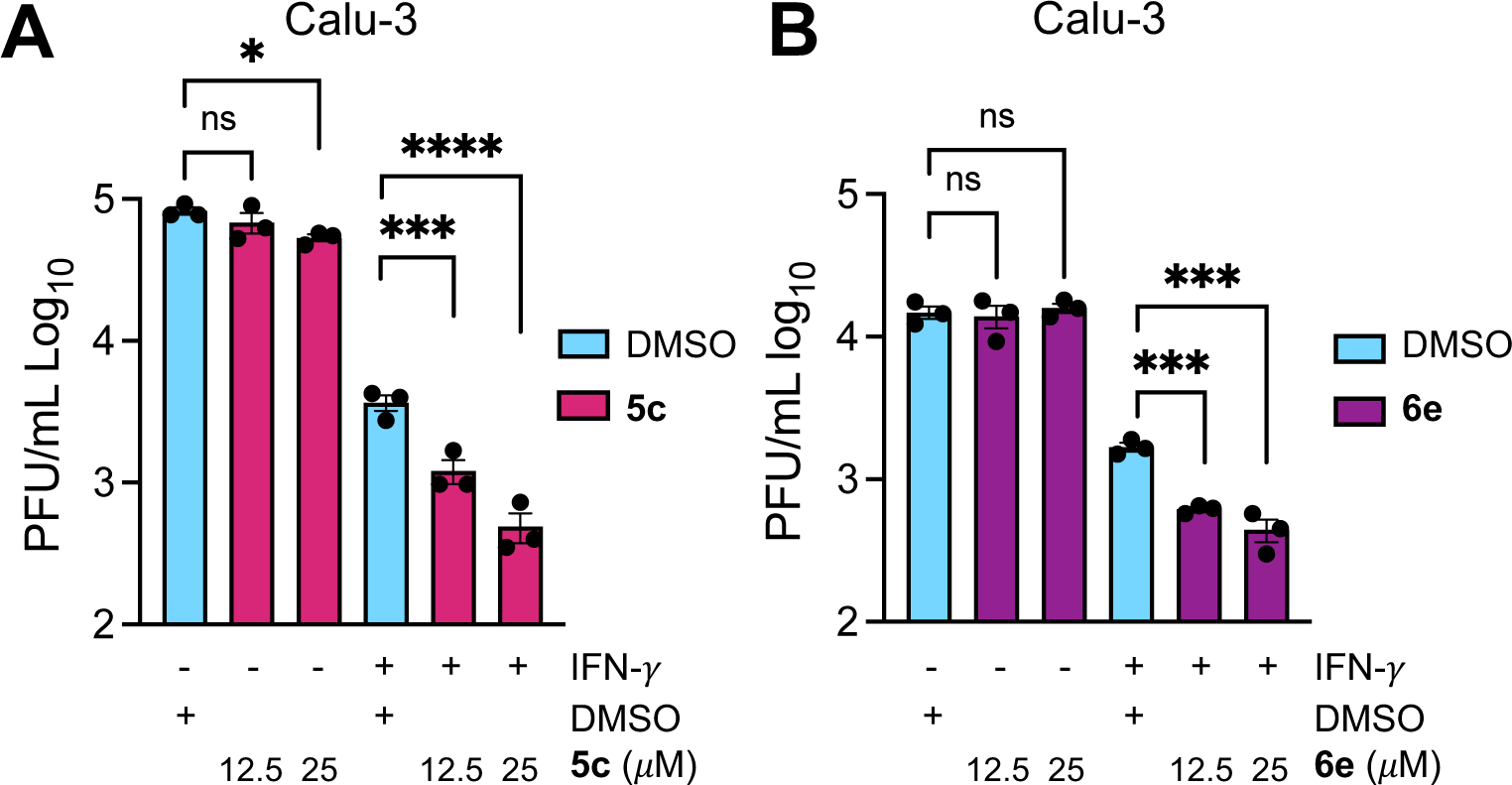
Compounds 5c and 6e inhibit SARS-CoV-2 replication. A-B) Calu-3 cells were mock or IFN-*γ* pre-treated (100 U) for 18 h, then were infected with SARS-CoV-2 and at 1 hpi the indicated concentration of **5c** (A) or **6e** (B) was added to the media. Cells and supernatants were collected at 20 hpi and progeny virus was measured by plaque assay. The results in A-B are from one experiment representative of 3 independent experiments. N=3 biological replicates.

To further demonstrate the specificity of our compounds, we looked to identify drug-resistant mutations in JHMV. We used JHMV for these experiments to avoid any potential gain of function issues with SARS-CoV-2 virus. We passaged 3 separate biological replicates of JHMV 3X in the presence of 150 *μ*M **5a** (5a1, 5a2, 5a3) (Fig. 8A). At this concentration **5a** inhibits infectious virus production by ∼0.5 logs, so that we would get a suitable concentration of virus produced to continue passaging (Fig. 8B). Virus exposed to **5a** became resistant by passage 2. We then took passage 3 virus and plaque-picked two separate biological replicates twice before sequencing the macrodomain from each isolate. In one of the plaque picked viruses, we identified two macrodomain mutations in neighboring residues, A1438T and G1439V (Fig. 8C). Remarkably, these are the same mutations that appeared in previous work with a different compound (40). Furthermore, we had engineered a recombinant virus with these mutations in a prior study evaluating different point mutants of Mac1, which demonstrated that this virus replicated at near WT levels in cell culture but was highly attenuated in mice (17). Notably, the A1438T/G1439V recombinant virus had increased replication compared to WT virus in the presence of **5a** (Fig. 8D). This result demonstrates that these mutations confer some resistance to **5a** and suggests that it targets Mac1. We also found that this virus had increased replication in the presence of **5c**, though the increase was not statistically significant (Fig. 8D).

**Fig 8.**
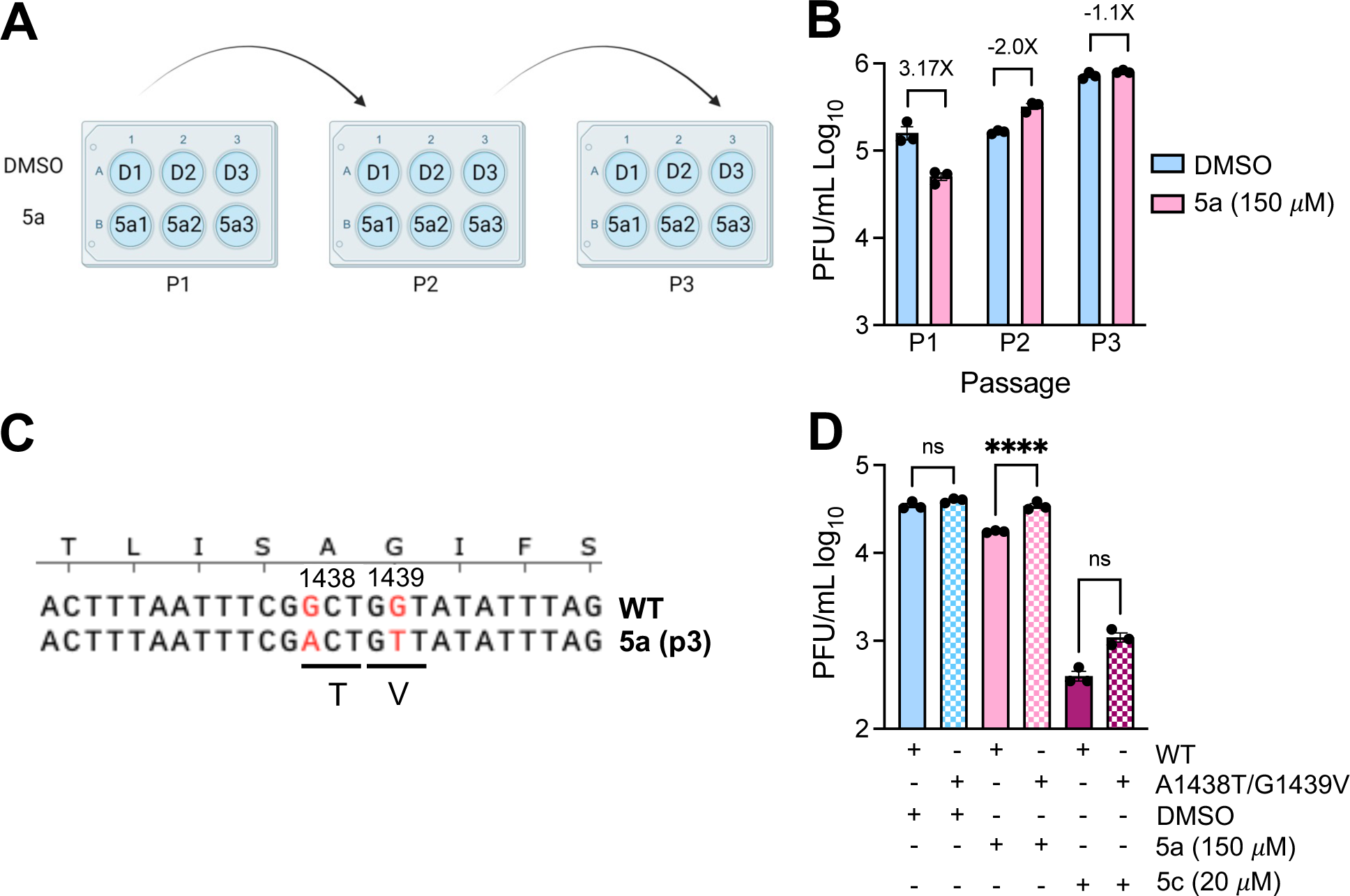
Identification of a Mac1 drug-resistant mutation. A) Cartoon depiction of passaging method for creating drug-resistant virus. 3 separate wells of DBT cells were initially infected with 0.1 MOI MHV in the presence of DMSO or **5a**. Each well was then passaged by taking 100 μl of cells/supernatants from the prior passage and infected a new well of DBT cells. Image was created using BioRender.com. B) Cells and supernatants were collected at 18-20 hpi at each passage and progeny virus was measured by plaque assay. C) Progeny virus at passage 3 was sequenced, which identified a two amino acid A1439T/G1439V mutation. D) DBT cells were infected with WT or A1438T/G1439V recombinant virus at an MOI of 0.1 in the presence of DMSO, **5a**, or **5c**. Cells and supernatants were collected at 20 hpi and progeny virus was measured by plaque assay. The data in D is from 1 experiment representative of 3 independent experiments. N=3 biological replicates.

In total, we developed a series of pyrrolo-pyrimidine based compounds that inhibit Mac1 activity *in vitro* and also repress both MHV and SARS-CoV-2 replication in cell culture by specifically targeting Mac1.

## DISCUSSION

Coronaviruses (CoVs) remain a global health threat, as demonstrated by the SARS-CoV-2 pandemic and earlier outbreaks of SARS-CoV and MERS-CoV. Over the past two decades, research on the conserved CoV macrodomain (Mac1) has found that it is critical for pathogenesis and promotes virus replication in the presence of interferon (IFN). The development of Mac1 inhibitors offers therapeutic potential and serves as a valuable strategy for probing the underlying mechanisms by which Mac1 promotes viral pathogenesis. In this study, we focused on improving previously developed Mac1 inhibitors as both chemical and antiviral agents (39).

While reverse genetics has proven to be a powerful tool in understanding Mac1 biology, there are several limitations to CoV reverse genetic systems. First, they are not available to all researchers; deletion mutations are not always recoverable and may have undesired impacts on neighboring genes, point mutations may not fully attenuate the functions of the protein, and they do not allow for temporal evaluation of function. Developing molecular probes targeting SARS-CoV-2 Mac1 would help uncover Mac1’s biological functions and advance our understanding of CoV biology, particularly how CoVs evade the host immune response. Specifically, Mac1 targeting probes offer an additional method to investigate how ADP-ribosylation, a process reversed by Mac1, impacts the interaction between the virus and host immune responses and disease outcomes during CoV infection both *in vitro* and *in vivo*. This is exemplified by previous studies where SARS-CoV-2 Mac1 deletion or mutation in animal models leads to attenuation of viral replication and enhanced interferon production, suggesting that effective Mac1 inhibition could weaken the virus while strengthening host defenses (18, 19).

Previously we expanded upon a prior fragment screen and identified several pyrrolo-pyrimidine based compounds with IC_50_ values less than 25 *μ*M. Pyrrolo-pyrimidine-based compounds are promising candidates as Mac1 inhibitors due to their molecular mimicry of adenine, which enables them to fit effectively into the ADP-ribose binding pocket of Mac1. This mimicry facilitates strong interactions within the Mac1’s active site, making these compounds valuable starting points for inhibitor development (35, 39). The most potent pyrrolo-pyrimidine from our previous work was **4a** (MCD-628), a tryptophanate. Furthermore, its enantiomer, **4b**, had even more potent inhibitory activity against Mac1 with an IC_50_ below 2 *μ*M. Here, we solved the crystal structure of **4a** with Mac1, which revealed key interactions such as hydrogen bonds with the amino acids D22 and I23, nearby water molecules and between the indole NH group and L126. While **4a** inhibited Mac1 activity *in vitro*, its physicochemical properties, most notably a prominent carboxylic acid that contributed to its negative logD value, were significant impediments to its antiviral activity. Many of the published Mac1 inhibitors, including those we synthesized, contain polar or acidic moieties that could limit their cellular permeability, which is a key determinant in translating *in vitro* activity into cell culture and *in vivo* efficacy. To address this problem, those moieties were initially modified to methyl and isopropyl ester groups to improve the lipophilicity of the compounds, as demonstrated with derivatives **5a** and **5c**, which had substantially improved logD values compared to **4a**. Despite the modest impact of these modifications on Mac1 inhibition, increasing the logD value correlated with their ability to inhibit virus replication. Interestingly, the **4a**, but not the **4b**-derived esters, inhibited Mac1 activity *in vitro* despite **4b** being the more potent inhibitor. Based on the co-crystal structure of **4a** and the docking model of **4b** (Fig. 2) the esterification could create some steric clash with the pyrimidine in the active conformation binding to Mac1. This was also observed in amide derivatives **6a-6e** where only **6e** derived from **4b** was active and predicted to have a distinct binding mode from the enantiomer **6d** (Fig. 6E).

To further explore the potential for replacing the carboxylate of **4a**/**4b** with more hydrophobic molecules, we introduced several amide-coupled pyridines (series **6**). None of these modifications substantially improved the IC_50_ of this series, as their IC_50_ values ranged from 4.04 μM (**6c**) to 8.37 μM (**6a**). Despite their ability to inhibit Mac1 *in vitro*, **6a-6c** were unable to repress virus replication, while **6d** and **6e** were modest inhibitors of MHV replication, and **6e** also inhibited SARS-CoV-2 in the presence of IFN-*γ*. The reason for this discrepancy is unclear, though it is noted that both **6d** and **6e** have nitrogen atoms in the 2 position of the pyridine ring, while **6a** and **6b** have the nitrogen in the 4 position. Regardless, structural insights from inhibitors such as **4a** and **4b** provide a foundation for developing next-generation inhibitors that can more effectively enter cells and demonstrate robust antiviral activity *in vivo*. Future strategies to optimize the Mac1 inhibitor antiviral activity will include *i)* improving their pharmacokinetic properties, *ii)* developing alternative delivery methods such as using nanoparticles to address the drug-delivery challenges, and *iii)* structural modifications that enhance the compounds’ ability to penetrate deeper into the Mac1 binding pocket and increase binding affinity.

To demonstrate the specificity of our hit compounds for Mac1 during infection, we first tested both **5c** and **6e** for inhibition of SARS-CoV-2 in the presence and absence of IFN-*γ*, as we have done previously (40). Each compound only inhibited virus replication in the presence of IFN-*γ*, which strongly indicates that these compounds target Mac1, as it is unlikely that there are other viral proteins where inhibition would demonstrate such stark differences between IFN-*γ*-treated and untreated cells. To further confirm specificity, we passaged MHV in the presence of **5a** with the goal of identifying drug-resistant mutations. We identified a resistant virus, and interestingly, it contained a two-amino-acid mutation in Mac1, A1438T/G1439V, which we observed previously after passaging MHV in the presence of a separate compound (40). Having the same mutation appear after passaging MHV in the presence of two different Mac1 inhibitors indicates that Mac1 is highly constrained in the ADP-ribose binding pocket and has limited options for developing resistance. We also previously reported that MHV A1438T/G1439V is highly attenuated in mice, indicating a fitness trade-off with the development of drug resistance against Mac1 targeting compounds.

In summary, our findings underscore the potential of pyrrolo-pyrimidine-based compounds as both therapeutic agents and molecular probes, enabling more profound insights into the role of Mac1 in CoV biology. By refining these inhibitors through targeted structural modifications and addressing pharmacokinetic limitations, we will continue to enhance their efficacy and therapeutic profiles. As we further investigate the mechanisms of Mac1 and its impact on viral replication, we will enable the development of innovative strategies to mitigate the threat posed by highly pathogenic human coronaviruses and potentially other emerging pathogens.

## METHODS

### Chemistry

See supplemental methods.

### Cell culture and reagents

Delayed brain tumor (DBT), L929, Vero E6, HeLa cells expressing the MHV receptor carcinoembryonic antigen-related cell adhesion molecule 1 (CEACAM1) (HeLa-MHVR), and baby hamster kidney cells expressing CEACAM1 (BHK-MVR) (all cell lines gifts provided by Stanley Perlman, University of Iowa) were grown in Dulbecco’s modified Eagle medium (DMEM) supplemented with 10% fetal bovine serum (FBS), 100 U/ml penicillin, 100 mg/ml streptomycin, HEPES, sodium pyruvate, nonessential amino acids, and L-glutamine. Calu-3 cells (ATCC) were grown in MEM supplemented with 20% FBS. Human IFN-*γ* was purchased from R&D Systems. ADP-ribosylated and control peptides were purchased from Cambridge peptides. Recombinant SARS-CoV-2 proteins was expressed with an N-terminal His-tag from a pET21a+ expression vector and purified as previously described (11).

### Crystallization, data collection, processing and refinement

Protein crystallization was performed using sitting-drop vapour-diffusion method in a Swissci 3D 96-well plate. The well solution contained the reported crystallization condition (33) but varied in PEG 3000 concentration: 0.1 M CHES pH 9.5, 28-32% PEG (v/v) 3000. 100 nL of 0.8 mM SARS-CoV-2 Mac1 was mixed with 100 nL of the crystallization solution using Mosquito pipetting robot (TTP Labtech). Crystals grew within 24 hr. Crystals from one droplet were then crushed, diluted in 100 µL reservoir solution and used as seeding solution. 0.8 mM SARS-CoV-2 Mac1 and 5 mM inhibitor were mixed and incubated at RT for 30 min. Crystallization drops were set up by mixing 100 nL of SARS-CoV-2 Mac1 and ligand solution and 100 nL of the reservoir solution. Then streak seeding was performed. Crystallization plates were monitored at RT with IceBear (47) and co-crystals appeared within 24 hr. For cryo-cooling, the crystals were soaked in 0.1 M CHES pH 9.5, 32% PEG (v/v) 3000 with 0.5 mM of the compound. X-ray diffraction data were collected on beamline BioMAX at MAX IV, Lund, Sweden. The dataset was processed by the XDS program package via XDSGUI (Table S1) (48, 49).

The structures were solved with PHASER (50) by the method of molecular replacement by using SARS-CoV-2 Mac1 (PDB: 8TV6) as a search model. Model building and refinement were performed with Coot (51) and REFMAC5 (52), respectively (Table S1). The structures were visualized in PyMOL version 1.7.2.1 (Schrödinger).

### Docking

The solved structure of 4a with Mac1 was prepared using the Schrödinger Protein Preparation Wizard (schrödinger.com), which adds hydrogens, predicts protonation status of titratable groups, optimizes hydrogen bonds, and then performs a constrained minimization. Only the water networks near the ligand were retained. Ligands were prepared using LigPrep and then docked into the receptor using Glide with XP precision (53, 54). The top scoring models were refined using Prime mmGBSA, allowing flexibility of the ligand and any residue/water within 5 Å of the ligand (55, 56).

### AlphaScreen (AS) assay

The AlphaScreen reactions were carried out in 384-well plates (Alphaplate, PerkinElmer, Waltham, MA) in a total volume of 40 μL in buffer containing 25 mM HEPES (pH 7.4), 100 mM NaCl, 0.5 mM TCEP, 0.1% BSA, and 0.05% CHAPS. All reagents were prepared as 4× stocks and 10 μL volume of each reagent was added to a final volume of 40 μL. All compounds were transferred acoustically using ECHO 555 (Beckman Inc) and preincubated after mixing with purified His-tagged macrodomain protein (250 nM) for 30 min at RT, followed by addition of a 10 amino acid biotinylated and ADP-ribosylated peptide [ARTK(Bio) QTARK (Aoa-RADP)S] (Cambridge peptides) (625 nM). After 1 h incubation at RT, streptavidin-coated donor beads (7.5 μg/mL) and nickel chelate acceptor beads (7.5 μg/mL); (PerkinElmer AlphaScreen Histidine Detection Kit) were added under low light conditions, and plates were shaken at 400 rpm for 60 min at RT protected from light. Plates were kept covered and protected from light at all steps and read on BioTek plate reader using an AlphaScreen 680 excitation/570 emission filter set. For counter screening of the compounds, 25 nM biotinylated and hexahistidine-tagged linker peptide (Bn-His6) (PerkinElmer) was added to the compounds, followed by addition of beads as described above. For data analysis, the percent inhibition was normalized to positive (DMSO + labeled peptide) and negative (DMSO + macrodomain + peptide, no ADPr) controls. The IC_50_ values were calculated via four-parametric non-linear regression analysis constraining bottom (=0), top (=100), & Hillslope (=1) for all curves.

### Differential scanning fluorimetry (DSF)

Thermal shift assay with DSF involved use of LightCycler® 480 Instrument (Roche Diagnostics). In total, a 15 μL mixture containing 8× SYPRO Orange (Invitrogen), and 10 μM macrodomain protein in buffer containing 20 mM HEPES-NaOH, pH 7.5 and various concentrations of ADP-ribose or hit compounds were mixed on ice in 384-well PCR plate (Roche). Fluorescent signals were measured from 25 to 95℃ in 0.2 C/ 30/Sec steps (excitation, 470–505 nm; detection, 540–700 nm). The main measurements were carried out in triplicate. Data evaluation and Tm determination involved use of the Roche LightCycler® 480 Protein Melting Analysis software, and data fitting calculations involved the use of single site binding curve analysis on GraphPad Prism. The thermal shift (ΔT_m_) was calculated by subtracting the T_m_ values of the DMSO from the T_m_ values of compounds. **Determination of LogD.** 0.11 mg of each compound weighed out into 2 mL centrifuge tubes. 1 mL of octanol saturated PBS or 0.1M HCl was added to each centrifuge tube and vortexed to dissolve. Then, 0.5-1.0 mL of the PBS saturated octanol was added to the 0.5-1.0 mL octanol saturated PBS supernatant and vortexed. Next, 100 *μ*l of the octanol saturated PBS phase from each centrifuge tube was added to separate UPLC vials. 100 *μ*l of 50:50 MP H_2_O:ACN was added to each vial and vortexed. The remaining octanol saturated PBS layer was removed from the vial using a micropipette and stored in separate vial. 100 *μ*l of the PBS saturated octanol layer from each centrifuge tube was added to separate UPLC vials. 100 *μ*l of 50:50 MP H_2_O:ACN was added to each vial and vortexed. These procedures were repeated in triplicate and then analyzed by UPLC/UV-VIS.

### Cell viability assay

Delayed brain tumor (DBT), L929 and Calu3 Cellular metabolic activity was assessed using a CyQUANT MTT cell proliferation assay (Thermo Fisher Scientific) by following the manufacturer’s instructions.

### Generation of recombinant pBAC-JHM constructs

Recombinant pBAC-JHMV^IA^ constructs were created using Red recombination as previously described (15). For pBAC-JHMV^IA^-nLuc, the nano-luciferase gene was amplified with ends homologous to the 5’ and 3’ end of ORF4 using the following primers:

F 5’-GGCAGCAAGTAGTTATGGCCCTCATCGGTCCCAAGACTACTATTGCTGCT GTCTTCACACTCGAAGATTTCG-3’ R 5’-GGCGTCACTCACAAGCCAAATCTCCATGTAGCTGGTGG TTACGCCAGAATGCGTTCGCACAGCCGCCAGCCGGTCA GCCAGTGTTACAACCAATTAAC-3’

The PCR product was recombined into pBAC-JHMV^IA^ replacing the ORF4 gene, creating pBAC-JHMV^IA^-nLuc (Fig. S1). pBAC-JHMV^IA^-A1438T/G1439V was previously described (17). BAC DNA was analyzed by restriction enzyme digest, PCR, and direct sequencing for isolation of correct clones.

### Reconstitution of recombinant pBAC-JHMV-derived virus

Approximately 1 × 10^6^ BHK-MVR cells were transfected with approximately 0.5 to 1 μg of pBAC-JHMV^IA^ DNA and 1 μg of pcDNA-MHV-N plasmid using PolyJet (SignaGen) as a transfection reagent. Stocks of the resulting virus were created by infecting ∼1.5 × 10^7^ 17Cl-1 cells at a multiplicity of infection (MOI) of 0.1 PFU/cell and collecting both the cells and supernatant at 16 to 20 hpi. The cells were freeze-thawed, and debris was removed prior to collecting virus stocks. Virus stocks were quantified by plaque assay on HeLa-MHVR cells and sequenced with the collection of infected 17Cl-1s or DBT cells using TRIzol. RNA was isolated and cDNA was prepared using MMLV-reverse transcriptase per the manufacturer’s instructions (Thermo Fisher Scientific). The nLuc gene sequence was amplified by PCR using the same primers as described above for sequencing BACs, and then resulting PCR products were sequenced by Sanger sequencing. The sequence was analyzed using DNA Star software.

### Virus infection

DBT and L929 were infected at an MOI of 0.1 with MHV strains JHMV^IA^-WT or JHMV^IA^-nLuc. Calu-3 cells were infected at an MOI of 0.1 with recombinant SARS-CoV-2 (Wuhan strain). For Calu-3 cells, trypsin-TPCK (1 μg/mL) was added to the medium at the time of infection. All infections included a 1-h adsorption phase, except for Calu-3 cells where the adsorption phase was increased to 2 h. Compounds were added after the adsorption phase. Infected cells and supernatants were collected at indicated time points and titers were determined on Hela-MHVR (MHV) or Vero E6 cells (SARS-CoV-2). For IFN-*γ* pretreatment experiments, human IFN-*γ* was added to Calu-3 cells 18 to 20 h prior to infection and were maintained in culture media throughout the infection.

### Identification of drug resistant MHV mutant viruses

DBT cells were infected in triplicate as described above. After each infection, the viral titer from each individual well was determined and then passaged to a new well of DBT cells. After 3 passages, 2 consecutive plaque picks were performed from 2 of the 3 individually passaged viral samples to collect individual isolates of MHV. RNA was isolated from these isolates using Trizol per manufacturer’s instructions. cDNA was prepared using MMLV-reverse transcriptase per the manufacturer’s instructions (Thermo Fisher Scientific) and PCR was performed using the following primers: Forward 5’-ggctgttgtggatggcaagca-3’ and Reverse 5’-gctttggtaccagcaacggag-3’. PCR products were sequenced by Sanger Sequencing (Azenta).

### Statistical analysis

All statistical analyses were done using a multivariant t-test to assess differences in mean values between groups, and graphs are expressed as geometric means ± geometric standard deviations (SD) (virus titers) or ± standard errors of the means (SEM). All data were analyzed using GraphPad Prism software. Significant *p* values are denoted with asterisks: *, *p* ≤ 0.05; **, *p* ≤ 0.01; ***, *p* ≤ 0.001.

### Data availability

Atomic coordinates and structure factors will be available at the Protein Data Bank with the id. 9GUB. Raw diffraction data will be available at fairdata.fi (https://doi.org/10.23729/c2152e19-38ec-4092-878d-f353358cbe5a).

## ACKNOWLEDGEMENTS

We thank Stanley Perlman for cell lines, Michael Hageman and the KU BIO Center for performing ADME studies, Mr. Kristopher Mason and Vaccitech for the use of their mass spectrometer, the Oulu Structural Biology core facility, a member of Biocenter Finland, Instruct-ERIC Centre Finland and FINStruct. ARF would like to thank funding from the NIH (R35GM138029), the NIH-funded Chemical Biology of Infectious Diseases (CBID) COBRE at the University of Kansas (P20GM113117), a CTSA grant from NCATS awarded to the University of Kansas for Frontiers: University of Kansas Clinical and Translational Science Institute (#UL1TR002366), a J.R. and Inez Jay Award from the University of Kansas, and a graduate student fellowship from the University of Kansas Madison and Lila Self graduate fellowship (JJP). DF would like to thank funding from the McDaniel College Student-Faculty Summer Research Fund, the Jean Richards Fund, the Schofield fund, and the Scott and Natalie Dahne fund. LL would like to thank funding from the Sidrid Jusélius foundation.

The funders had no role in study design, data collection and analysis, decision to publish, or preparation of the manuscript.

## Author contributions

Conceptualization: JJP, LL, DVF, ARF

Data curation: JJP, MTHD, DC, DKJ, NS, AR, LL, DVF, ARF

Formal analysis: JJP, MTHD, DC, DKJ, AR, LL, DVF, ARF

Funding acquisition: JJP, LL, DVF, ARF

Methodology: JJP, MTHD, DKJ, AR, LL, DVF, ARF

Investigation: JJP, MTHD, DC, LMS, IC, GC, DT, JP, NS, JJOC, PS, DKJ, AR, LL, DVF, ARF

Project administration: LL, DVF, ARF

Resources: DKJ, AR, LL, DVF, ARF

Visualization: JJP, MTHD, DC, LMS, IC, GC, DT, JP, NS, DKJ, AR, LL, DVF, ARF

Validation: JJP, MTHD, DKJ, AR, LL, DVF, ARF

Supervision: DKJ, AR, LL, DVF, ARF

Writing—original draft: JJP, ARF

Writing—review & editing: JJP, MTHD, DC, LMS, IC, GC, DT, JP, NS, JJOC, PS, DKJ, AR, LL, DVF, ARF

A.R.F. was named as an inventor on a patent filed by the University of Kansas for a live-attenuated SARS-CoV-2 vaccine.

