## Supplemental Files for "Identification of a series of pyrrolo-pyrimidine based SARS-CoV-2 Mac1 inhibitors that repress coronavirus replication"

**CONTENT**

**SUPPLEMENTAL METHODS**
**FIGURES S1-S4**
**TABLE S1**

### SUPPLEMENTAL METHODS

#### Chemistry Section

The chloride **1** was protected with tosyl chloride to afford the protected compound **2** in 83% yield. Nucleophilic aromatic substitution was conducted on chloride **2** with either D- or L-tryptophan methyl ester to afford the enantiomers **3a** (*S*) and **3b** (*R*) in good yield. Hydrolysis of the tosyl group and the methyl ester was done in methanolic sodium hydroxide to afford the enantiomers **4a** (*S*) and **4b** (*R*). The carboxylic acids **4a** and **4b** were esterified under typical Fischer esterification conditions to afford the methyl esters **5a** (*S*) and **5b** (*R*) and isopropyl esters **5c** (*S*) and **5d** (*R*). Amide couplings were also conducted with carboxylates **4a** and **4b** to afford amides **6a-e**.

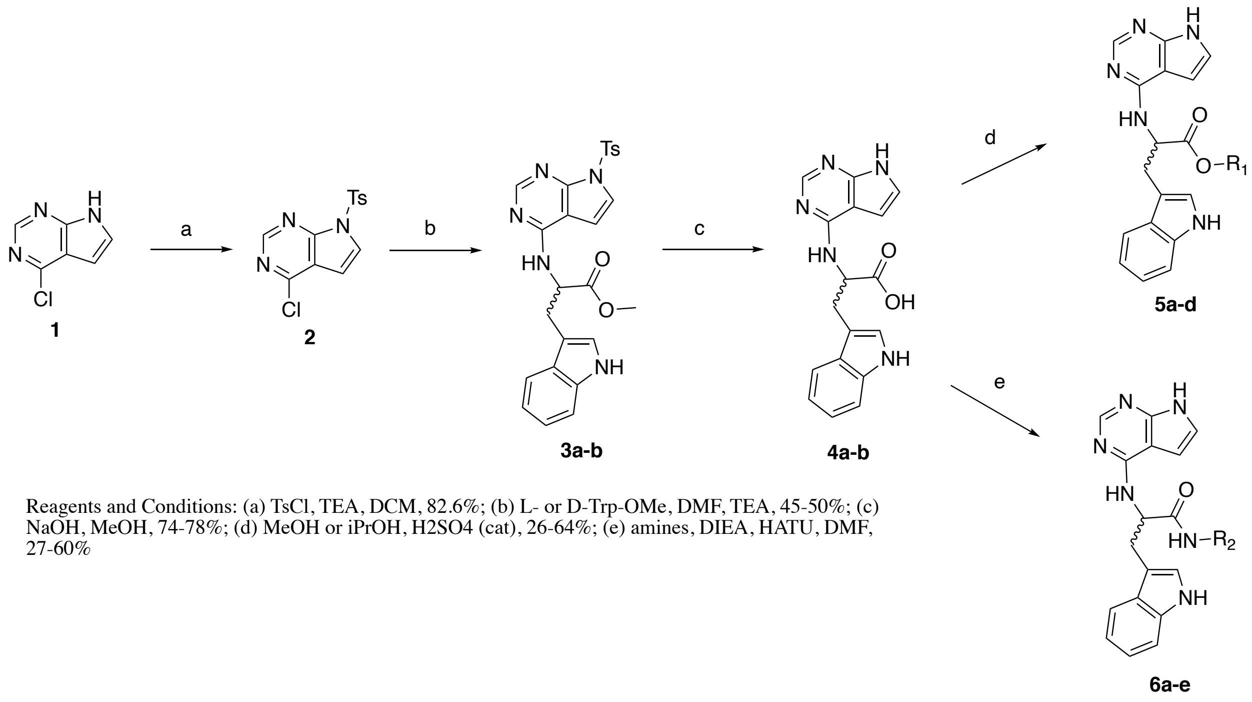


Reagents and Conditions: (a) TsCl, TEA, DCM, 82.6%; (b) L- or D-Trp-OMe, DMF, TEA, 45-50%; (c) NaOH, MeOH, 74-78%; (d) MeOH or iPrOH, H2SO4 (cat), 26-64%; (e) amines, DIEA, HATU, DMF, 27-60%

**Scheme 1. Synthesis of 4-substituted 7*H*-pyrrolo[2,3-*d*]pyrimidine derivatives.**

#### Experimental Section

All solvents were reagent grade or high-performance liquid chromatography (HPLC) grade. Unless otherwise noted, all materials were obtained from commercial suppliers and used without further purification. $^{1}H$ NMR spectra were recorded at 400.19 MHz. All $^{13}C$ spectra were recorded at 100.63 MHz. The HPLC solvent system consisted of distilled water and acetonitrile, both containing 0.1% formic acid. Chiral HPLC was conducted using a Diacel Chiralpak® AD-3 column (150 x 4.6mm i.d., 3mm), solvent system hexane/ethanol/DEA/TFA – 80/20/0.1/0.1. All final compounds tested were confirmed to be of ≥95% purity by the HPLC methods described above. HRMS were run in positive ion mode on a SCIEX X500B QTOF mass spectrometer coupled to an Exion LC. Each compound was resuspended in 0.1% formic acid and injected directly on to the mass spec by flow injection.

**(MCD-666) 4-chloro-7-tosyl-7*H*-pyrrolo[2,3-*d*]pyrimidine (2)**. The 4-chloro-7*H*-pyrrolo[2,3-*d*]pyrimidine (10g, 65mmol) and triethyl amine (2 equiv) were dissolved in 50 mL of DCM. A solution of the tosylate (12.4g, 65 mmol) in DCM (50 mL) was added dropwise over the course of an hour under nitrogen. The reaction was stirred overnight and then quenched with water (50 mL). The reaction was partitioned and the DCM layer was washed once more with water (50 mL). The DCM layer was then separated, dried with $MgSO_4$, filtered and concentrated in vacuo. The resulting solid was triturated twice with boiling hexanes (~100mL) and the resulting tan solid was isolated and characterized as the desired material. Dry yield = 16.5g (82.6%). $^1$HNMR ($CDCl_3$): 8.77 (s, 1H), 8.09 (d, 2H, $J$ = 8.4 Hz), 7.78 (d, 1H, $J$ = 4.0 Hz), 7.32 (d, 2H, $J$ = 8.4 Hz), 6.71 (d, 1H, $J$ = 4.0 Hz), 2.40 (s, 3H). $^{13}$CNMR ($CDCl_3$): 153.14, 152.56, 151.00, 146.31, 134.39, 129.99, 128.35, 127.28, 119.80, 103.00, 21.68; HRMS (ESI+): *m/z* calculated for $C_{13}H_{10}ClN_3O_2S$ (M+1)+ 308.0182, Found 308.0241.

**(MCD-677) Methyl (7-tosyl-7*H*-pyrrolo[2,3-*d*]pyrimidin-4-yl)-*L*-tryptophanate (3a). Methyl (7-tosyl-7*H*-pyrrolo[2,3-*d*]pyrimidin-4-yl)-*L*-tryptophanate (3a).** To a 100 mL sealed round bottom flask, the 4-chloro-7-tosyl-7*H*-pyrrolo[2,3-*d*]pyrimidine (3.3 g, 10.7 mmol), methyl *L*-tryptophanate (3.0 g, 11.8 mmol), TEA (5.0 mL, equiv), and DMF (10 mL) were added and stirred at just below reflux overnight. The DMF was removed using a rotary evaporator and was redissolved in ethyl acetate (~25 mL) and water (~25 mL). The organic layer was then further purified using column chromatography using 50/50 ethyl acetate/hexanes. The final yield was 4.1 g (76.6 %). $^1$HNMR ($CDCl_3$): 8.46 (s, 1H), 8.17 (s, 1H), 8.03 (d, 2H, $J$ = 8.0 Hz), 7.47 (d, 1H, $J$ = 7.6 Hz), 7.42 (d, 1H, $J$ = 4.0 Hz), 7.35 (d, 1H, $J$ = 7.6 Hz), 7.27 (d, 2H, $J$ = 8.0 Hz), 7.18 (t, 1H, $J$ = 7.6 Hz), 7.05 (t, 1H, $J$ = 7.6 Hz), 6.99 (s, 1H), 6.25 (d, 1H, $J$ = 4.0 Hz), 5.5 (bs, 1H), 5.27 (m, 1H), 3.70 (s, 3H), 3.44 (m, 2H), 2.37 (s, 3H). $^{13}$CNMR ($CDCl_3$): 172.72, 155.30, 153.68, 150.54, 145.44, 136.18, 135.20, 129.71, 128.09, 127.67, 122.82, 122.71, 122.35, 119.78, 118.47, 111.34, 109.99, 105.40, 101.52, 54.04, 52.34, 27.71, 21.61. HRMS (ESI+): *m/z* calculated for $C_{25}H_{23}N_5O_4S$ (M+1)+ 490.1471, Found 490.1515.

**(MCD-893) Methyl (7-tosyl-7*H*-pyrrolo[2,3-*d*]pyrimidin-4-yl)-*D*-tryptophanate (3b).** This compound was made in an identical manner to **3a** with identical spectroscopic data. Dry yield = 4.3 g (82.2%). HRMS (ESI+): *m/z* calculated for $C_{25}H_{23}N_5O_4S$ (M+1)+ 490.1471, Found 490.1533.

**(MCD-628) (*7H*-pyrrolo[2,3-*d*]pyrimidin-4-yl)-*L*-tryptophan (4a).** The protected ester **3a** (250 mg, 0.51 mmol) was dissolved in MeOH (1 mL) followed by addition of 10% NaOH (1 mL, xs) and heating to reflux for 3 hours. The solvent was then removed in vacuo and the crude residue was acidified to pH 4.0 while stirring and cooling. The resulting solid was filtered off and triturated with cold water (~5mL) and filtered off again. The resulting solid was characterized as the desired compound **4a**. Dry yield = 121mg (74%). $^1$HNMR (DMSO-$d_6$): 11.46 (s, 1H), 10.75 (s, 1H), 8.01 (s, 1H). 7.57 (d, 1H, $J$ = 8.0Hz), 7.50 (s, 1H), 7.28 (d, 1H, $J$ = 8.0Hz), 7.16 (s, 1H), 7.01 (m, 3H), 6.58 (s, 1H), 4.82 (m, 1H), 3.22 (m, 2H); $^{13}$CNMR (DMSO-$d_6$): 174.52, 155.62, 151.06, 150.17, 136.05, 127.19, 123.60, 120.96, 120.83, 118.31, 118.19, 111.34, 110.60, 102.59, 98.70, 54.19, 27.23; Chiral HPLC – 7.29 min (>98% enantiomeric excess). HRMS (ESI+): *m/z* calculated for $C_{17}H_{15}N_5O_2$ (M+1)+ 322.1226, found 322.1262.

**(MCD-709) (*7H*-pyrrolo[2,3-*d*]pyrimidin-4-yl)-*D*-tryptophan (4b).** This compound was made in an identical manner to **4a** with identical spectroscopic properties. Dry yield = 192 mg (78%). Chiral HPLC – 5.76 min (>98% enantiomeric excess). HRMS (ESI+): *m/z* calculated for $C_{17}H_{15}N_5O_2$ (M+1)+ 322.1226, found 322.1271.

**(MCD-617) methyl (7*H*-pyrrolo[2,3-*d*]pyrimidin-4-yl)-*L*-tryptophanate (5a).** The carboxylic acid **4a** (200mg, 0.62mmol) was dissolved in methanol (5 mL) followed by the addition of 5 drops of $H_2SO_4$ and refluxing overnight. The methanol was removed followed by partitioning between 10% $NaHCO_3$ and EtOAc (10 mL each). The organic layer was dried with $MgSO_4$, filtered and concentrated to afford an amorphous solid that was triturated with DCM (1 mL) and hexanes (1 mL). The dry yield was 0.115g (55%). $^1$HNMR ($CDCl_3$): 10.24 (br s, 1H), 8.38 (s, 1H), 8.16 (br s, 1H), 7.57 (d, 1H), 7.35 (d, 1H), 7.17 (t, 1H), 7.09 (t, 1H), 7.02 (m, 2H), 6.24 (d, 1H), 5.61 (d, 1H), 5.39 (d, 1H), 3.70 (s, 3H), 3.49 (m, 2H). $^{13}$C NMR (DMSO-$d_6$): 173.58, 155.37, 151.01, 150.23, 136.06, 127.04, 123.69, 121.15, 120.93, 118.40, 118.00, 111.43, 110.06, 102.58, 98.69, 54.02, 51.72, 27.27; HRMS (ESI+): *m/z* calculated for $C_{18}H_{17}N_5O_2$ (M+1)+ 336.1382, found 336.1418.

**(MCD-888) methyl (7*H*-pyrrolo[2,3-*d*]pyrimidin-4-yl)-*D*-tryptophanate (5b).** This compound was made in an identical manner to **5a** using **4b** (200mg, 0.62 mmol) as the starting material. The dry yield was 0.135g (64.9%). Spectroscopic properties were identical to **5a**. HRMS (ESI+): *m/z* calculated for $C_{18}H_{17}N_5O_2$ (M+1)+ 336.1382, found 336.1423.

**(MCD-814) isopropyl (7*H*-pyrrolo[2,3-*d*]pyrimidin-4-yl)-*L*-tryptophanate (5c).** The carboxylic acid **4a** (200mg, 0.62mmol) was dissolved in isopropanol (10 mL) followed by addition of three drops of conc. sulfuric acid. The reaction was heated for 24 hours after which the TLC indicated that the reaction was complete. The isopropanol was removed in vacuo and the resulting residue was partitioned between EtOAc (10 mL) and 10% $NaHCO_3$ (5 mL). The organic layer was washed with $NaHCO_3$ once more, dried with $MgSO_4$, filtered and concentrated to afford a foam. The foam was triturated with DCM/hexanes to afford 60 mg of the desired ester (26.6%). $^1$HNMR (DMSO-$d_6$): 11.49 (br s, 1H), 10.79 (br s, 1H), 8.00 (s, 1H), 7.64 (d, 1H, $J$ = 7.6 Hz), 7.54 (d, 1H, $J$ = 7.2 Hz), 7.29 (d, 1H, $J$ = 7.6 Hz), 7.19 (s, 1H), 6.97-7.07 (m, 3H), 6.62 (s, 1H), 4.83 (m, 2H), 3.22 (m, 2H), 1.13 (d, 3H, $J$ = 6.4 Hz), 0.94 (d, 3H, $J$ = 6.4 Hz). $^{13}$CNMR ($CDCl_3$): 173.17, 155.91, 151.42, 150.76, 136.57, 127.62, 124.18, 121.55, 121.37, 118.82, 118.53, 111.85, 110.51, 103.07, 99.21, 67.98, 55.05, 27.79, 21.97, 21.68. HRMS (ESI+): *m/z* calculated for $C_{20}H_{21}N_5O_2$ (M+1)+ 364.1695, found 364.1751.

**(MCD-875) isopropyl (7*H*-pyrrolo[2,3-*d*]pyrimidin-4-yl)-*D*-tryptophanate (5d).** This compound was made in an identical manner to **5c** using **4b** (200mg, 0.62 mmol) as the starting material. The dry yield was 0.124g (55%). Spectroscopic properties were identical to **5a**. HRMS (ESI+): *m/z* calculated for $C_{20}H_{21}N_5O_2$ (M+1)+ 364.1695, found 364.1749.

**(MCD-710) (*S*)-2-((7*H*-pyrrolo[2,3-*d*]pyrimidin-4-yl)amino)-3-(1*H*-indol-3-yl)-*N*-(pyridin-4-ylmethyl)propenamide (6a).** To a 25 mL round bottom flask, carboxylic acid **4a** (100 mg, 0.31mmol) was equipped with a magnetic stir bar. HATU (174.9 mg, 0.46 mmol), DIEA (0.1 mL, 0.62mmol), and approximately 2 mL of DMF were added to the flask and stirred overnight. TLC (EtOAc) indicated that the reaction was complete. The DMF was removed in vacuo and the crude product was partitioned between ethyl acetate (10 mL) and 10% sodium bicarbonate (5 mL). The organic layer was dried with $MgSO_4$, filtered and concentrated in vacuo. Methanol/ethyl acetate (1/1) was used to triturate the product which was collected by vacuum filtration. Dry yield: 47mg (36.8%). $^1$HNMR (DMSO-$d_6$): 11.47 (br s, 1H), 10.77 (br s, 1H), 8.70 (t, 1H, $J$ = 5.6 Hz), 8.38 (d, 2H, $J$ = 6.0 Hz), 8.08 (s, 1H), 7.70 (d, 1H, $J$ = 7.6 Hz), 7.51 (d, 1H, $J$ = 8.0 Hz), 7.28 (d, 1H, $J$ = 8.0 Hz), 7.22 (d, 1H, $J$ = 2.4 Hz), 7.14 (d, 1H, $J$ = 5.6 Hz), 6.96-7.06 (m, 5H), 6.65 (d, 1H $J$ = 2.4 Hz), 4.98 (m, 1H), 4.29 (m, 2H), 3.21 (m, 2H). $^{13}$CNMR ($CDCl_3$): 173.50, 156.11, 151.53, 150.69, 149.71, 149.05, 136.54, 127.72, 124.29, 122.31, 121.39, 121.32, 118.99, 118.70, 111.76, 111.07,

103.25, 99.39, 55.55, 41.56, 28.39. HRMS (ESI+): *m/z* calculated for $C_{23}H_{21}N_7O$ (M+1)+ 412.1808, found 412.1861.

**(MCD-770) (*S*)-2-((7*H*-pyrrolo[2,3-*d*]pyrimidin-4-yl)amino)-*N*-((2-chloropyridin-4-yl)methyl)-3-(1*H*-indol-3-yl)propenamide (6b).** To a 25 mL round bottom flask, carboxylate **4a** (100 mg, 0.31mmol) (2-chloropyridin-4-yl)methanamine (66.24 mg, 0.37 mmol), HATU (174.9 mg, 0.46 mmol), DIEA (0.1 mL, 0.62mmol), and approximately 2 mL of DMF were stirred overnight. TLC plate showed the reaction was complete (EtOAc). The DMF was removed in vacuo and the crude product was partitioned between ethyl acetate (10 mL) and 10% sodium bicarbonate (5 mL). The organic layer was dried with $MgSO_4$, filtered and concentrated in vacuo. Ethyl acetate was used to triturate the product which was collected by vacuum filtration. Dry yield: 94mg (60.0%). $^1$HNMR (DMSO-$d_6$): 11.47 (br s, 1H), 10.76 (br s, 1H), 8.74 (s, 1H), 8.24 (d, 1H, $J$ = 4.4 Hz), 8.12 (s, 1H), 7.71 (d, 1H, $J$ = 8.0 Hz), 7.54 (d, 1H, $J$ = 6.8 Hz), 7.18-7.33 (m, 4H), 6.97-7.04 (m, 3H), 6.64 (s, 1H), 4.91 (m, 1H), 4.35 (m, 2H), 3.19 (m, 2H). $^{13}$CNMR (DMSO-$d_6$): 173.74, 156.13, 153.56, 151.57, 150.81, 150.67, 149.90, 136.54, 127.66, 124.30, 122.37, 121.79, 121.43, 121.33, 118.92, 118.71, 111.80, 111.07, 103.27, 99.37, 60.18, 55.73, 28.25. HRMS (ESI+): *m/z* calculated for $C_{23}H_{20}ClN_7O$ (M+1)+ 446.1418, found 446.1480.

**(MCD-804) (*S*)-2-((7*H*-pyrrolo[2,3-*d*]pyrimidin-4-yl)amino)-3-(1*H*-indol-3-yl)propenamide (6c).** To a 25 mL round bottom flask, **4a** (200 mg, 0.62mmol), ammonium chloride (0.04g, 0.74 mmol), HATU (353 mg, 0.93 mmol), DIEA (0.32 mL, 1.86mmol), and approximately 2 mL of DMF were stirred overnight. The DMF was removed in vacuo and the crude product was partitioned between ethyl acetate (10 mL) and 10% sodium bicarbonate (5 mL). The organic layer was dried with $MgSO_4$, filtered and concentrated in vacuo. Ethyl acetate was used to triturate the product which was collected by vacuum filtration. Dry yield: 55 mg, (27%). $^1$HNMR (DMSO-$d_6$): 11.44 (br s, 1H), 10.72 (br s, 1H), 8.01 (s, 1H), 7.68 (d, 1H, $J$ = 7.6 Hz), 7.51 (s, 1H), 7.34 (d, 1H, $J$ = 6.8 Hz), 7.25 (d, 1H, $J$ = 7.6 Hz), 7.16 (d, 1H, $J$ = 2.0 Hz), 6.92-7.02 (m, 4H), 6.60 (d, 1H, $J$ = 2.0 Hz), 4.92 (m, 1H), 3.20 (m, 2H). $^{13}$CNMR (DMSO-$d_6$): 175.02, 156.10, 151.55, 150.65, 136.47, 127.81, 124.01, 121.30, 121.22, 119.01, 118.62, 111.68, 111.33, 103.12, 99.28, 54.87, 28.36. HRMS (ESI+): *m/z* calculated for $C_{17}H_{16}N_6O$ (M+1)+ 321.1386, found 321.1438.

**(MCD-808) (*S*)-2-((7*H*-pyrrolo[2,3-*d*]pyrimidin-4-yl)amino)-3-(1*H*-indol-3-yl)-*N*-(pyridin-2-ylmethyl)propenamide (6d).** The reaction was conducted in a similar manner to **6a**. Dry yield = 48mg (39%). $^1$HNMR (DMSO-$d_6$): 11.46 (br s, 1H), 10.74 (br s, 1H), 8.71 (t, 1H, $J$ = 6.0 Hz), 8.43 (d, 1H, $J$ = 4.8 Hz), 8.07 (s, 1H), 7.70 (d, 1H, $J$ = 7.6 Hz), 7.65 (t, 1H, $J$ = 7.6 Hz), 7.49 (s, 1H), 7.29 (d, 1H, $J$ = 7.6 Hz) 7.04-7.22 (m, 3H), 6.93-7.03 (m, 3H), 6.63 (s, 1H), 5.00 (m, 1H), 4.35 (m, 2H), 3.16 (m, 2H). $^{13}$CNMR ($CDCl_3$): 173.40, 159.18, 156.12, 151.54, 150.68, 149.08, 136.94, 136.52, 127.75, 124.26, 122.31, 121.38, 121.29, 121.03, 119.00, 118.68, 111.73, 111.15, 103.23, 99.37, 55.52, 44.66, 28.36. HRMS (ESI+): *m/z* calculated for $C_{23}H_{21}N_7O$ (M+1)+ 412.1808, found 412.1842.

**(MCD-809) (*R*)-2-((7*H*-pyrrolo[2,3-*d*]pyrimidin-4-yl)amino)-3-(1*H*-indol-3-yl)-*N*-(pyridin-2-ylmethyl)propenamide (6e).** This compound was made in a similar manner to **6d** using the carboxylate **4b** as the starting material. The product had identical spectroscopic data. Dry yield = 55mg (43%). HRMS (ESI+): *m/z* calculated for $C_{23}H_{21}N_7O$ (M+1)+ 412.1808, found 412.1844.

### SUPPLEMENTAL FIGURES

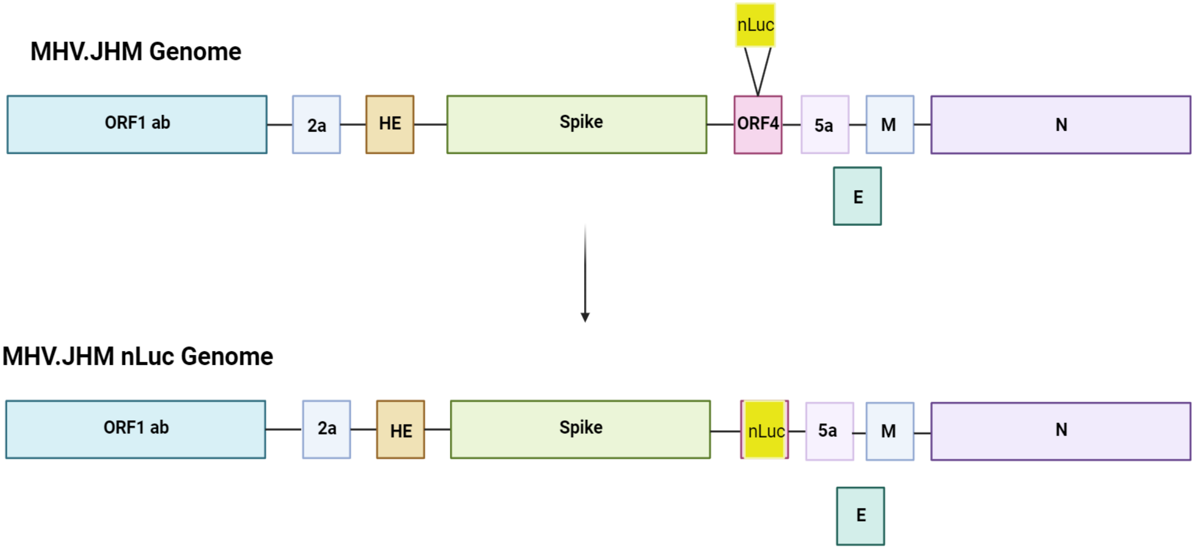


**Fig. S1. Creation of an MHV-JHM expressing nanoluciferase.** The nanoluciferase gene was inserted into the MHV genome in place of ORF4 using lambda-red recombination.

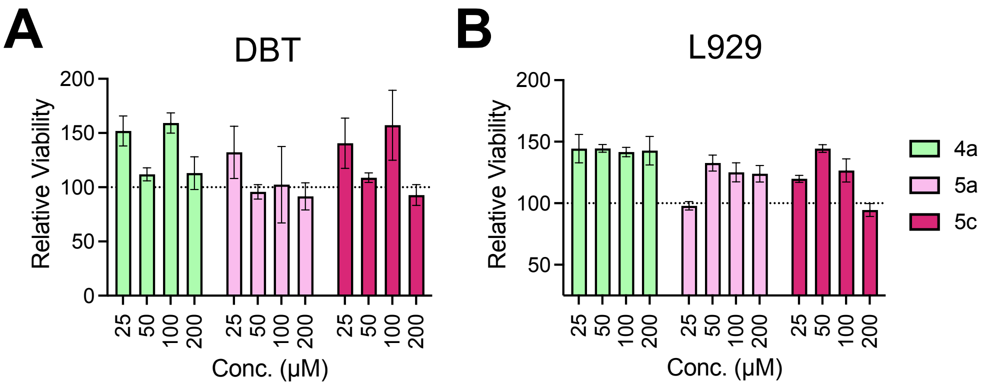


**Fig. S2. Compounds 4a, 5a, and 5c do not impact the viability of cells used to measure MHV replication.** A-D) Cell viability for compounds **4a**, **5a**, and **5c** (A-B) on DBT (A) and L929 (B) cells was measured using an MTT assay. The data in A-B are from one experiment representative of 3 independent experiments.

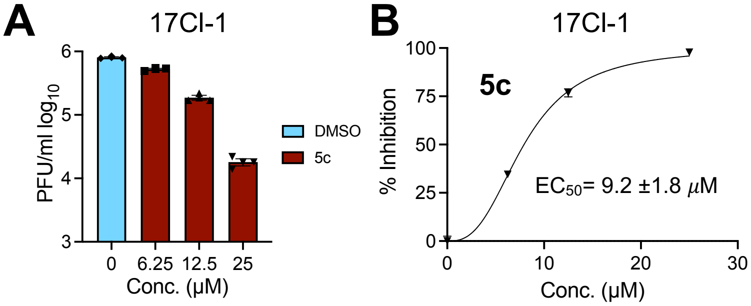


**Fig. S3. Compound 5c inhibits MHV replication on 17Cl-1 cells.** A-B) 17Cl-1 cells were infected with JHMV-WT and at 1 hpi the indicated concentration of **5c** was added to the media. Cells and supernatants were collected at 20 hpi and progeny virus was measured by plaque assay. The results are from 1 experiment representative of 2 independent experiments. n=3 biological replicates. The $EC_{50}$ value was averaged over 2 independent experiments and includes standard deviation.

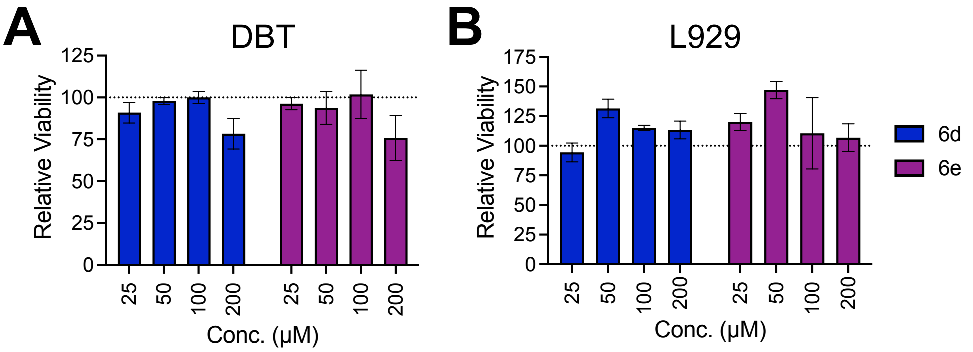


**Fig. S4. Compounds 6d and 6e do not impact the viability of cells used to measure MHV replication.** A-B) Cell viability for compounds **6d** and **6e** (A-B) on DBT (A) and L929 (B) cells was measured using an MTT assay. The data in A-B are from one experiment representative of 3 independent experiments.

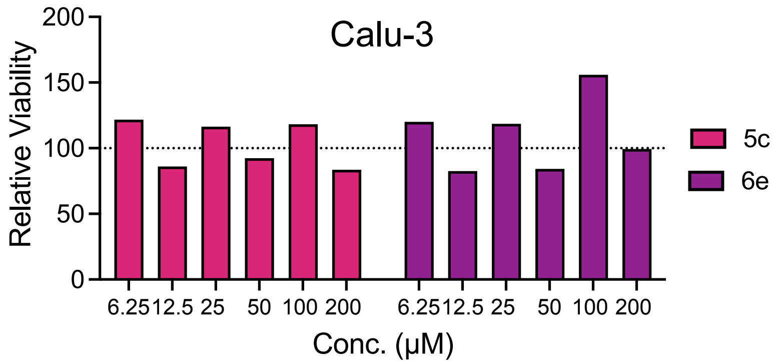


**Fig. S5. Compounds 5c and 6e do not impact the viability of Calu-3 cells used for SARS-CoV-2 infections.** Cell viability for compounds 5c and 6e on Calu-3 cells was measured using an MTT assay. The data are from one experiment representative of 3 independent experiments.

### SUPPLEMENTAL TABLE

**Table S1.** Data collection and refinement statistics for Mac1-**4a** co-crystal structure.

| **PDB id.** | **9GUB** |
|---|---|
| **Data collection** | |
| Beamline | MAX IV, BioMAX |
| Wavelength (Å) | 0.8266 |
| Space group | $P1\ 2_1\ 1$ |
| Unit cell dimensions | |
| *a, b, c* (Å) | 37.53, 32.85, 120.6 |
| α, β, γ (°) | 90, 94.738, 90 |
| Resolution range (Å) | 40.06 - 1.10 (1.13 - 1.10) |
| Total no. of reflections | 805989 (59194) |
| No. of unique reflections | 119450 (8807) |
| Completeness (%) | 100% (100%) |
| $\langle I/\sigma(I)\rangle$ | 15.34 (1.99) |
| CC1/2 (%) | 100 (71.3) |
| $R_{meas}$ | 5.6 (103.7) |
| **Model building and refinement** | |
| R-factor | 15.52 |
| R-free | 17.48 |
| No. of atoms | |
| Protein | 2654 |
| Ligands | 48 |
| Water | 347 |
| RMSD | |
| Bonds (Å) | 0.009 |
| Angles (°) | 1.501 |
| Average *B* factors ($Å^2$) | |
| Protein | 17.66 |
| Ligands | 20.66 |
| Water | 27.99 |
| Ramachandran plot | |
| Favoured (%) | 98.17 |
| Allowed (%) | 1.83 |
| Outliers (%) | 0.0 |

[1]Values within parentheses refers to the highest resolution shell.
